# Performance of a Modular Robotic Cluster Matches Skilled Human Operators for Complex Cell Therapy Manufacturing Tasks

**DOI:** 10.1101/2025.04.25.650551

**Authors:** Sudeshna Sadhu, Brigitte Schmittlein, Angela Lares, Christopher Cheng, Xiaojie Chen, Varun Bhatia, Winston Zha, Sebastien Wha, Sunaina Nayak, Marco Uboldi, Yifan Yu, Jiawei Wang, Wei Zhang, Prajakta P. Bhanap, Yongchang Ji, Andrew Scheffler, John Wilson, Dan Welch, Nikolaos Gkitsas-Long, Aidan Jeffrey Retherford, Ramya Tunuguntla, Alice Melocchi, Alexandre Girard, Federico Parietti, Steven A. Feldman, Jonathan H. Esensten

## Abstract

Although numerous cell and gene therapies have received regulatory approval, their adoption has been hampered by high cost and challenges in scaling out manufacturing. Many autologous cell therapies are individually manufactured for each patient using traditional manual methods in high-cost environments. Therefore, robotics and automation offer a potential solution to meet the growing demand for such therapies. To ensure identical biological outcomes, automation must replicate validated manual workflows, a requirement that poses significant engineering challenges, especially for aseptic manipulation and compatibility with manual-centric instruments and consumables. Here we describe the design and performance of a modular robotic cluster consisting of specialized modules containing widely adopted equipment. The robotic arm uses custom end-effectors to handle standard consumables and instruments such as syringes, vials, bags, cell counters, bioreactors, incubators, and closed centrifuges. We compared the performance of skilled human operators against the robotic cluster across multiple tasks: transferring cells between sterile bags, cell counting, drawing volume from a vial to a syringe, and resuspension and sampling from both a bag and G-Rex 100M-CS bioreactor. The robotic system also executed high-complexity operations with industry-standard instruments: cell selection using a CytoSinct 1000 and wash/buffer exchange with a CTS Rotea Counterflow centrifugation system. Experimental results for each unit operation show that the robotic cluster’s performance is equivalent to manual operations on the selected key metrics. These data demonstrate that the robotic system can efficiently and robustly perform specific unit operations, which can be combined in any order for end-to-end cell therapy manufacturing processes.

**One Sentence summary:** A modular robotic system can perform key unit operations in cell therapy manufacturing with accuracy comparable to human operators while ensuring throughput and flexibility.

## 1. Introduction

Cell therapy manufacturing is a complex process that involves isolating, modifying, expanding, and formulating living cells from a human donor for therapeutic use. Many of these therapies are autologous, which means that the same patient is the source and recipient of the therapeutic cells. Several autologous cell therapies have achieved regulatory approval for the treatment of cancer [1–6]. The number of clinical trials of cell therapies has dramatically increased: more than 3,000 trials were initiated over the last five years, with about 600 starting in 2023 alone [7–9]. The majority of them were sponsored by pharmaceutical companies, highlighting the commercial potential of these therapeutics. However, current manufacturing approaches are difficult to scale out and are unlikely to be able to satisfy patient demand [10–12]. Indeed, manufacturing of cell therapies in clean rooms requires multiple manual steps by highly trained operators [13,14]. This manual production strategy limits throughput and increases risk of errors, contamination, and manufacturing failures [10].

Despite rapid advances in cell therapy, automation efforts to date have been devoted to single production steps, while comprehensive end-to-end automation remains limited. Two major automation strategies have emerged: an all-in-one approach, in which multiple unit operations are performed by a single device, and a modular approach, in which instruments from different vendors perform specific unit operation in series [15–19]. All-in-one systems require skilled operators for setup, monitoring, and interventions, and they are harder to optimize and scale for new processes. Moreover, they generally lack standardized interfaces for connectivity and data exchange, limiting their use in combination with other instruments. On the other hand, the modular approach allows easier customization. Modular systems readily incorporate newly available equipment or instruments intended for specific processes and limit the reliance on a single supplier for any single task. Most importantly, modular workflows reduce the process constraints from the duration of the full process itself to the longest cycle-time of the isolated unit operations. This enables parallel processing and results in increased throughput capacity of each individual manufacturing line. Due to these advantages, a modular approach to cell therapy manufacturing has been extensively developed [20–26]. However, certain limitations for modular approach still persist: for instance this approach often requires manual intervention for transferring reagents/cells between different modules and instruments. These manual hand-offs introduce error and inconsistency, posing significant contamination risk and ergonomic challenges that can hinder both quality and throughput. Despite parallel processing advantages, manual interventions become a bottleneck, impacting rate and scalability of manufacturing. Relying on equipment from different vendors can also introduce compatibility and interoperability challenges and requires the need for custom interfaces for data integration and process control. Finally, open manual manipulations are still required in the modular approach for critical tasks such as sampling, media preparation, and quality control assays [20,24,27,28]. End-to-end robotic automation can overcome these bottlenecks and help unlock the broader clinical and commercial potential of cell-based therapies by improving robustness of modular manufacturing procedures and reducing costs [15–17]. Moreover, robotic automated systems enhance productivity per unit area, potentially decreasing labor costs.

As a preliminary step towards end-to-end automation, in this study we describe the development and performance of a modular robotic cluster capable of performing a variety of common cell therapy manufacturing unit operations in a flexible sequence. The robotic cluster includes multiple modular instruments within a single robotically automated platform. The cluster has a fully closed manufacturing workflow with minimized human intervention, thus reducing the risk of contamination. The cluster supports integration of hardware and software across multiple Good Manufacturing Practice (GMP)-validated instruments and consumables, all controlled from a single interface.

We have previously assessed the potential of this modular approach, building a proof-of-concept robotic cluster that used industry-standard equipment to successfully culture CD8+ T cells with comparable cell yields, viability, and identity to those generated through conventional manual processing [29]. We extend this work to demonstrate comparable performance between manual and robotic approaches for the following operations: bag-to-bag cell transfer, automated cell counting, drawing volume from a vial to a syringe, cell resuspension and sampling from both a bag and G-Rex 100M-CS bioreactor, cell selection using CytoSinct 1000 and wash/buffer exchange using CTS Rotea Counterflow centrifugation system. In each case, we compare the performance of the robot to human operators and establish the accuracy and robustness of the automated operations.

## 2. Results

### 2.1 The modular robotic cluster

The robotic cluster represents an evolution of the system previously described [29]. It consists of several modules, which are accessed by a robotic arm mounted on a central rail (Figure 1a). Each module contains standard equipment or consumables already used in GMP-compliant manufacturing processes. A combination of modules are used to replicate the manual cell therapy manufacturing process. The modules used in the present study are shown in Figure 1b in green. They have the following functions: the input/output module is for consumable addition and removal from the system. The consumable storage module enables intermediate storage during runs and vial-to-syringe fluid transfer operations. The cell expansion/counting module contains hardware necessary for reagent transfer, cell transfer between bags and bioreactors, and sampling. Cell selection using the CytoSinct 1000 magnetic cell separation system (GenScript) and cell harvest using CTS Rotea Counterflow centrifugation system (Thermo Fisher Scientific) were performed in the cell selection and counterflow centrifuge modules, respectively. The cell incubator module contains a cell culture incubator (Thermo Fisher Scientific, HERACELL VIOS 250i) capable of storing multiple G-Rex 100M-CS bioreactors (Cytiva) or cell culture bags.

**Figure 1.**
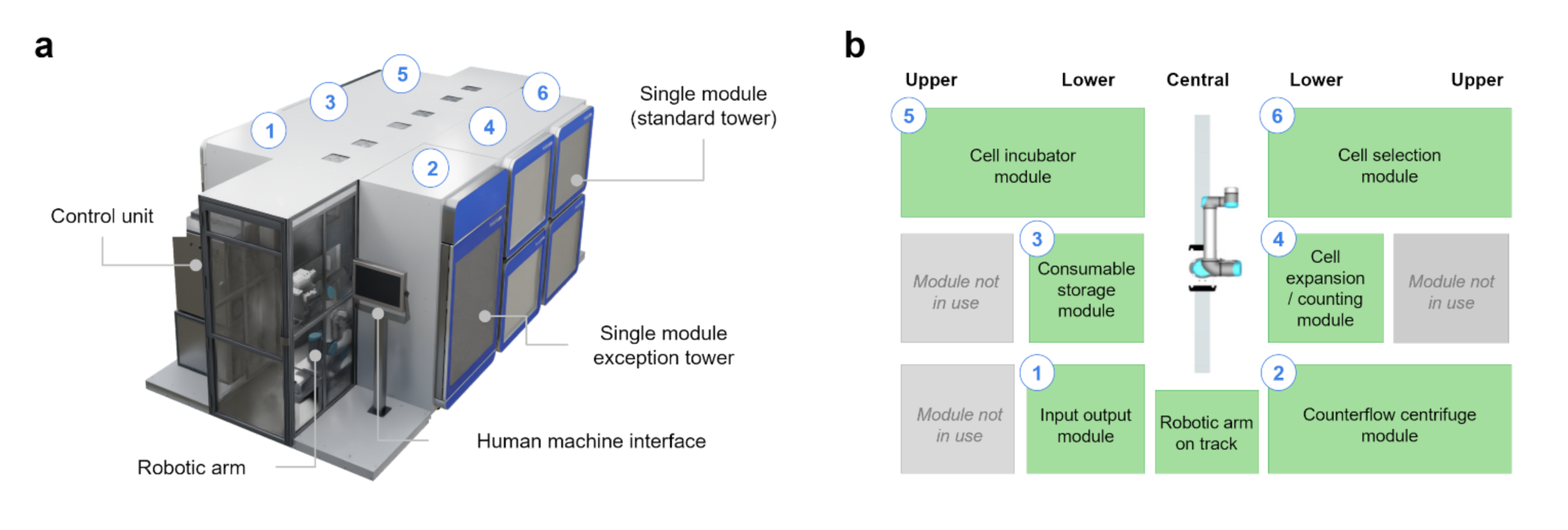
Schematics of the robotic cluster and modules. a) 3D rendered outline of the modular robotic cluster. b) Representative diagram (top view) of the different towers/modules used in this study.

The development of the robotic system’s hardware and software was guided by principles including unaltered reproduction of manual bioprocess, easy equipment/consumable validation against manual bioprocess, sterility, modularity, autonomy, traceability, robustness, and usability (Supplementary Material, Table S1). The core principle of our automation strategy is to precisely replicate the validated manual process. By replicating the manual manufacturing process, we can directly transfer a validated manual process to the automated system while minimizing or eliminating major process modifications and extensive comparability studies. The robotic system’s design directly implements this principle in two key ways. 1) The system is engineered to use commercially-available consumables (bags, vials, syringes or bioreactors) that are used in manual protocols, ensuring cells only contact materials already validated and the robotic manipulations are designed to emulate their manual counterparts to maintain equivalent process conditions (flow rate, shear stress, temperature, process timing). For some tasks, motions from a human operator were recorded and translated directly to the robot to ensure the system achieves a comparable or equivalent performance to a trained technician. 2) Another design principle is that the entire process is physically isolated from the external environment to minimize the risk of microbial contamination. Manual operations in a cleanroom rely on a variety of aseptic techniques such as working within a biosafety cabinet or using sterile welds for fluid transfer. With these manual techniques, prevention of microbial contamination is dependent on the skill of the operator. By contrast, the robotic system uses sanitized needle-free connectors for all fluidic transfers. The interior of the robotic cluster has an air filtering and monitoring system to further minimize contamination risk.

The robotic cluster hardware is designed with modular architecture to accommodate any combination of instruments based on the unique needs of the cell manufacturing process. Modularity is achieved through swapping modules with different instruments or functions, enabled by customizable module deckplates. There are minimum modifications performed to the off-the-shelf instruments and the associated consumable kits for easier and faster technology transfer process from manual to automated workflow. The robotic system software operates autonomously. The only human interaction with the robotic cluster involves loading and unloading consumables via the input/output module. To make manual interventions user-friendly, the cluster’s process planning and execution software (the “orchestrator”) allocates load and unload waves and displays detailed loading instructions in the user interface. For traceability, every operation performed in the robotic system, including reagent transfer volumes and instrument parameters, is tracked in the digital batch record. The orchestrator’s parallel planning algorithm improves manufacturing throughput by enabling a single robotic arm to switch between tasks for different batches, taking into account pre-defined biological time constraints.

### 2.2 Automated liquid handling approach

Closed connections between different vessels in traditional manual processes are typically done via tube welding. Tube welding is difficult to automate. It entails fitting flexible tubes into grooves on a metal plate, locking the tubes into place, pressing a button on the welding instrument, removing the tubes, and squeezing the weld site to open the interior fluid path. As an alternative for the robotic cluster, we developed rigid protected connectors [29].

Rigid connectors enable reliable connections and address all the disadvantages of tube welding. Operating rigid connectors based on a standardized thread (*e.g.* Luer lock) is straightforward for both human operators and robotic systems. A successful connection can be confirmed by measuring simple process parameters such as locking torque [Nm] or linear advancement along the thread [mm]. The connectors are designed to work together as a system. The sterile fluid path is closed unless connectors are mated to each other. Rigid connectors can be attached to known lengths of tubing and fixed to specified locations within the robotic cluster. This approach allows the robotic arm to find and utilize the connector at its fixed location.

One example of this approach is fluid transfer between a syringe and a vial (Figure 2). A customized port and coupler enable the robot to perform this process (Figure 2a,b). The port is a snap-on plastic adapter that surrounds a SmartSite needle-free female luer connector. The coupler is a snap-on plastic adapter that surrounds a Texium needle-free male luer connector. These proprietary adapters do not modify the connector and do not come in contact with the sterile fluid path. The port contains external features that allow the vial adapter to be docked in a fixed location. The port has conical lead-in features that enable the mating coupler to be reliably connected. The coupler has complementary internal lead-in features that guide in onto a Port and external conical features that allow the coupler end of arm tooling (EOAT) (Figure 2a,b) to grasp the coupler, self-center it within the gripper, and rotate it to engage the luer lock threads.

**Figure 2.**
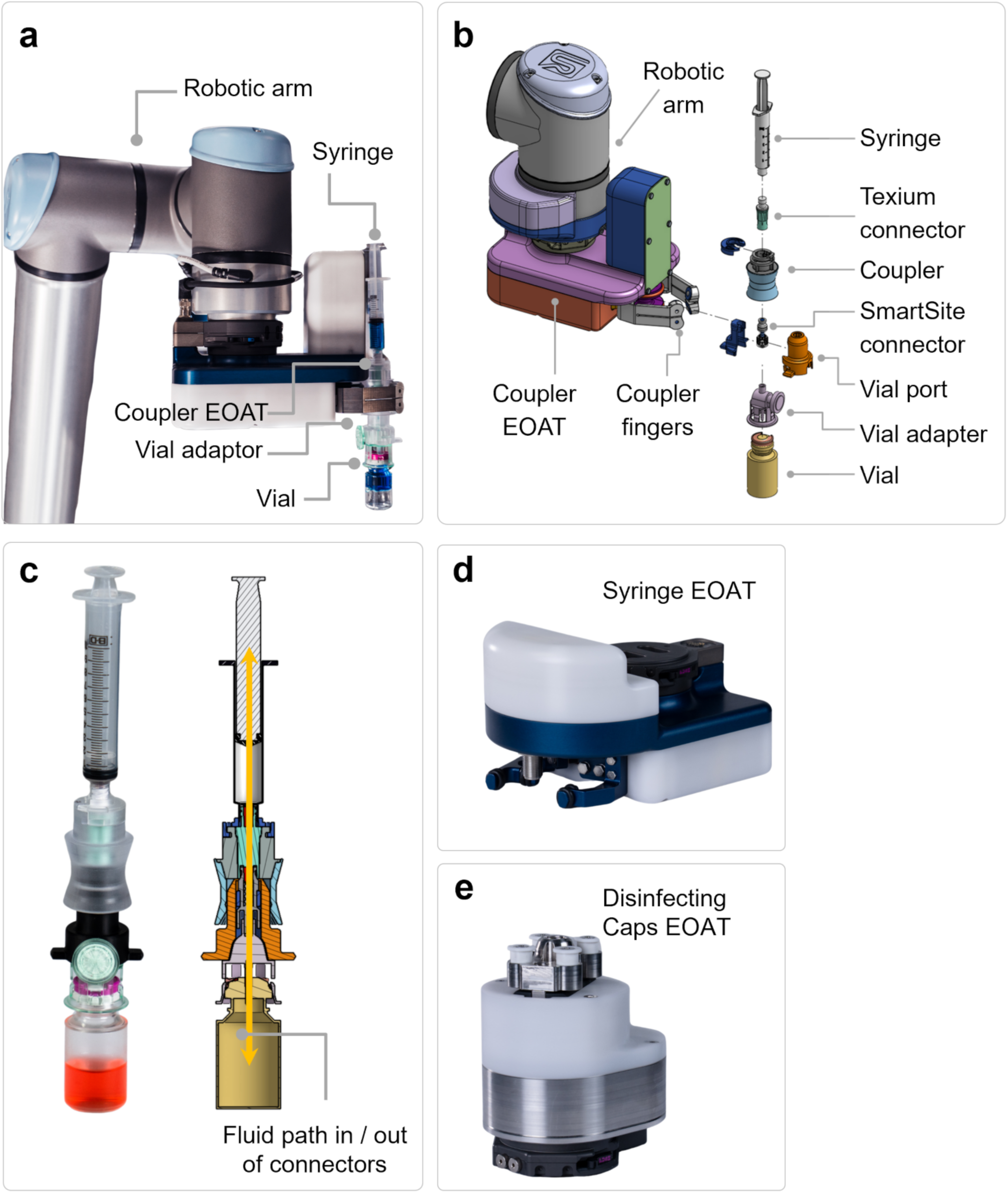
Liquid handling automation tools. (a) Image of the 6 axis robotic arm with a coupler EOAT grasping a vial and syringe assembly. (b) Exploded view diagram of the robotic vial and syringe assembly. (c) Section view for the connector assembly showing liquid flow path. (d) Image of the syringe EOAT. (e) Image of the disinfecting cap EOAT.

The robotic arm is equipped with a 6-axis force-torque sensor that enables it to estimate interaction forces and adjust positioning during the coupler-port mating interaction and ensure that the connection is centered. Monitoring for a proper connection is done via several sensors in the system. For example, the force-torque sensor in the robot arm checks that the expected force thresholds are met during the connection process. When the connector is fully engaged and bottoms out, force will spike in the vertical direction past a threshold.

For fluid transfer between a vial and a syringe, the port is attached to the vial adapter, and the coupler is attached to the syringe. When the syringe and vial are mated together, a pathway is opened for fluid to be transferred from the vial to the syringe (Figure 2c). The coupler EOAT grasps the vial and syringe assembly and allows the robot arm to invert the vial and slot its plunger into a syringe pulling fixture. This inversion allows fluid to be drawn from the vial into the syringe. After this step is completed, the vial is docked back into its cartridge, the syringe is decoupled, and then connected to the target vessel for dispensing. A syringe EOAT (Figure 2d) is connected to the robot arm and enables the arm to sense where the plunger of a syringe is, grasp it, and push or pull it to dispense or withdraw fluid (Supplementary Material, Movie S1). We designed robotic grippers to automatically operate disinfecting caps (Figure 2e). Disinfecting caps are commercially available, one-time-use components that contain a disinfecting solution and a thread that matches the shape of the SmartSite and Texium connectors. The use of disinfecting caps on the connectors before coupling further decreases the risk of microbial contamination.

### 2.3 Bag-to-bag cell transfer and vial-to-syringe volume draw

Withdrawal of liquid from a reagent vial with a syringe (Figure 3a) and bag-to-bag transfer of liquid (Figure 3b) are routine tasks in cell therapy manufacturing. The underlying design principles of the automated vial-to-syringe interaction were described above. The accuracy of a vial-to-syringe fluid draw was assessed using standard glass vials and 10 mL syringes. The consumable module in the robotic cluster includes a vial and syringe cartridge dock used for storage and manipulation of vials and syringes as well as a syringe puller. The robotic gripper picks the syringe coupler, places it on top of the vial and locks the vial and syringe port together by engaging the Texium connector onto the vial (Figure 3c). The vial-syringe assembly is then inverted and the barrel flange of the syringe is constrained to a fixed location on the syringe puller. The robotic gripper then pulls the syringe plunger to a specific distance for drawing the fluid.

**Figure 3:**
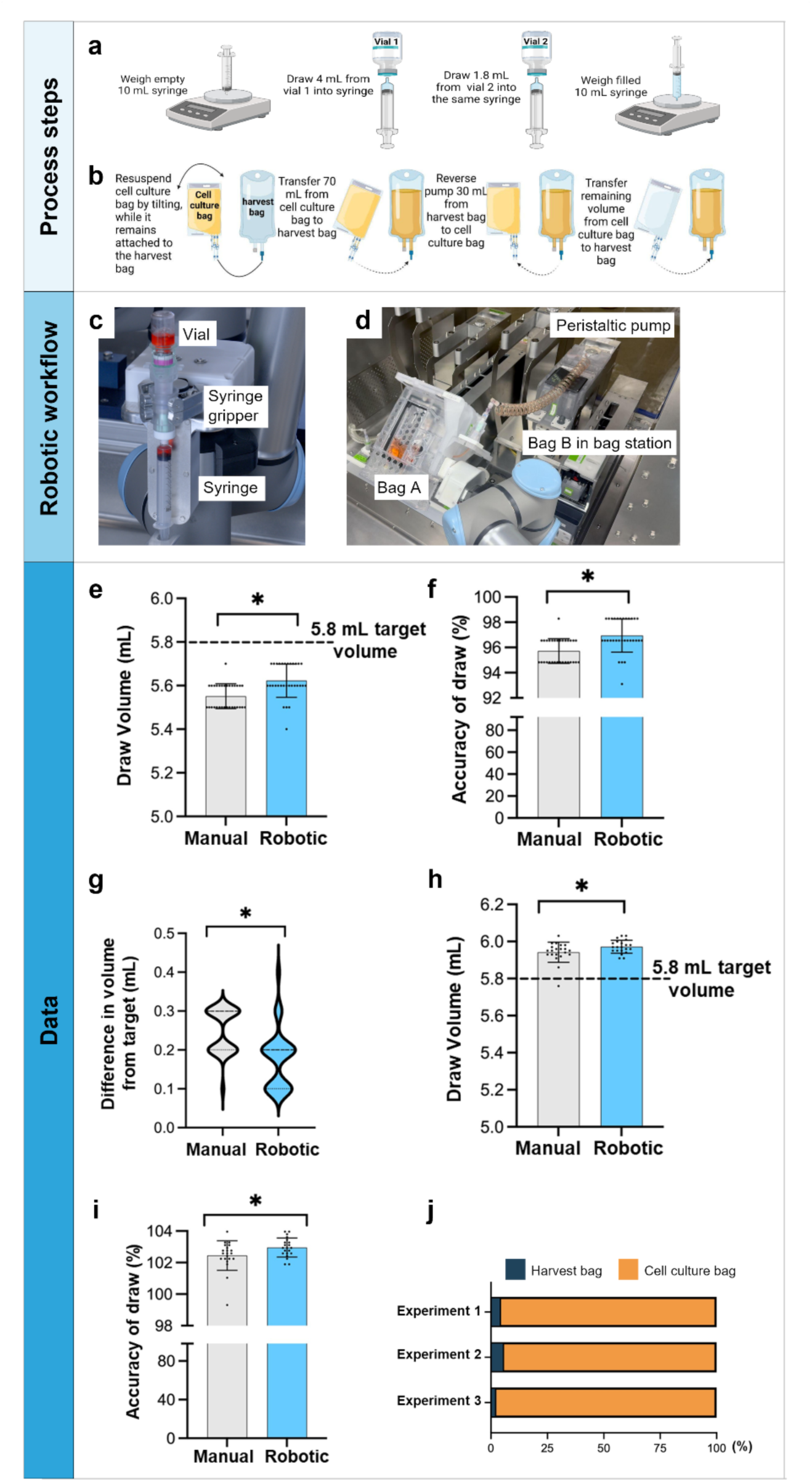
Accuracy of vial-to-syringe draw in manual and robotic conditions and automated bag-to-bag transfer of human T cells. (a-b) Schematics showing the robotic vial-to-syringe draw (a) and robotic bag-to-bag transfer (b) workflow. (c) Image showing robotic arm holding the vial to the syringe assembly in an inverted manner to draw fluid. (d) Transfer of cells from bag A to bag B using a peristaltic pump. (e-g) Data showing initial calibration conditions (unpaired t-test. n = 31). (h-i) Data showing modified calibration conditions (unpaired t-test, n = 20). (j) Data is represented as the percentage of cells in each bag after the transfer was complete. n = 3 independent experiments.* indicates statistically significant, p < 0.05.

For transferring liquid between two bags, we designed an in-house bag station consisting of load cells, peristaltic pumps and a dock for bag cartridges (Figure 3d). Off-the-shelf bags were custom-modified by attaching a coiled pump tube, ending in a Texium connector at one end and a SmartSite needle-free connector at the other. The modified bags were assembled on a bag cartridge compatible with robotic handling. The bag cartridge is aligned automatically with a peristaltic pump which allows fluid to be pumped to and from a bag in either direction. To achieve precise bag-to-bag liquid transfer, the software of the robotic cluster monitors the change in bag weight with load cells that were integrated within the bag station. Based on the change in bag weight, we computed the real time flow rate (change in bag weight over time) and dynamically adjusted the pump motor speed to accurately pump to the set target volume.

To compare side-by-side manual vs robotic vial-to-syringe draw accuracy, an arbitrary target volume of 5.8 mL was chosen as the total draw volume from two vials. The average draw volume was 5.55 mL for the manual condition and 5.62 mL for the robotic condition (Figure 3e). The accuracy was slightly higher in robotic draws compared to manual (Figure 3f). There was considerable overlap in the volume difference from the target volume between the manual and robotic conditions (Figure 3g). We calibrated the robotic system to account for dead volume in the syringe hub and performed an additional experiment. The target volume remained 5.8 mL. The average volume for the manual conditions was 5.94 mL and for the robotic condition 5.97 mL (Figure 3h). Both the manual and robotic conditions were able to achieve a draw volume slightly above the target volume (Figure 3i). To validate the robotic bag-to-bag transfer unit operation, we assessed the efficiency of cell transfer between bags. We cultured human T cells in 100 mL growth media in a source bag (bag A) and transferred the entire cell suspension to the final transfer bag (bag B) using a closed-loop pumping system with intermittent robotic resuspension of the source bag. Post robotic bag-to-bag transfer, we assessed the distribution of cells in both the source and the final bag. Across three experiments, an average of 96% of viable cells were successfully transferred, with 4% viable cells retained in the initial bag (Figure 3j).

### 2.4 Bioreactor or Bag Resuspension and Sampling

Cells tend to settle or clump together in the bags or bioreactors, leading to uneven distribution in the vessel. Mixing of the vessel uniformly distributes the cells, ensuring that subsequent samples are representative of the culture contents. In this unit operation, we manually or robotically mixed a bag or a bioreactor, sampled using a syringe, and measured the resulting cell concentration (Figure 4a-d). Bag mixing is automated through a simple robotic trajectory involving the 6th joint on the robot (Figure 4e). The trajectory is defined such that the robot’s tool center point rotates 180 degrees about the tool center point axis. In contrast, mixing of cells in a G-Rex bioreactor was automated by capturing an experienced scientist’s hand motions using the motion capture system OpiTrak. This motion was then extracted and converted into the base frame of the robot, enabling the robot to precisely mimic these movements when holding a G-Rex (Figure 4f). This approach can be extended to automate any custom mixing motions. To automate syringe sampling from a bag or bioreactor, the software communicates with a proximity sensor that is built into a custom end-effector of the robot. The software takes into account the conversion between robot tool center point movement in mm and syringe plunger displacement in mL to a precision of 0.1 mL. This approach allows the robot to accurately detect where the syringe plunger is and how much to pull to draw a particular volume of sample provided by the operator.

**Figure 4:**
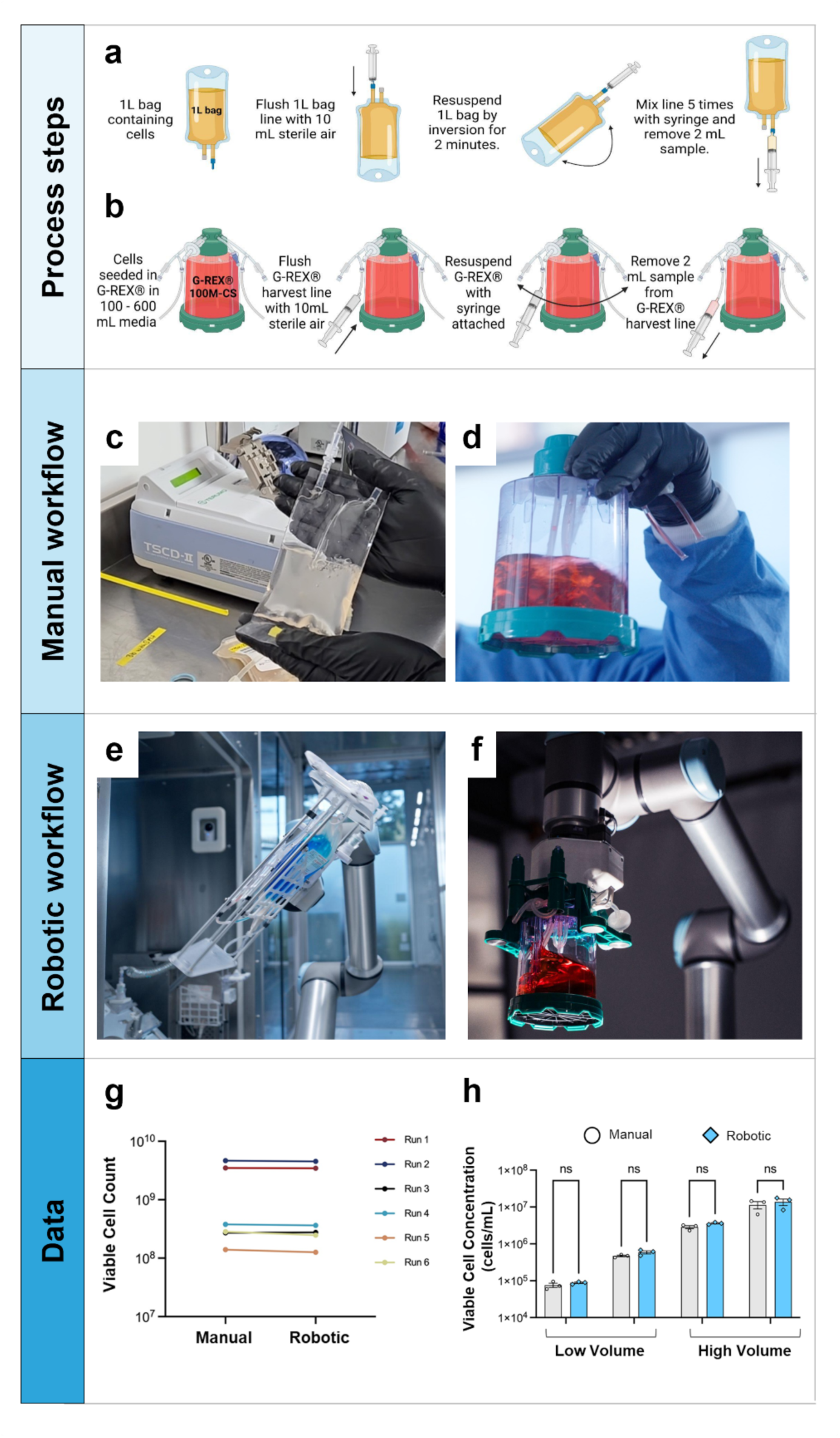
Manual versus robotic resuspension of cells in a bag and bioreactor. (a,b) Schematics showing robotic process steps for 1L transfer bag and G-Rex 100M-CS bioreactor resuspension and sampling respectively. (c,f) Image of bag and bioreactor resuspension performed manually (c,d) and robotically (e-f). (g,h) Manual versus robotic resuspension of human T cells in a 1L transfer bag and G-Rex 100M-CS bioreactor. Viable cell counts post resuspension were measured. Cells at different concentrations and volumes in a G-Rex 100M-CS bioreactor were resuspended with manual or robotic swirling motion (h). Cell concentration was then measured (unpaired t-test, n = 3).

We experimentally compared the performance of manual and robotic mixing processes. One liter transfer bags or a G-Rex 100M-CS bioreactor were assembled into proprietary robotic cartridges. Sampling was performed using either a 5 mL or 10 mL syringe assembled on a robotic coupler (Figure 4c,d). For comparing manual versus automated bag mixing, primary human T cells at low (1x10^6^ cells/mL) and high (20x10^6^ cells/mL) concentrations were placed in 50 - 300 mL volumes in 1L bags. The bags were mixed by manual or robotic mixing for 2 mins. The bags were then sampled using a syringe and counted (Figure 4a). On average, across six runs, the cell counts after robotic mixing were within 5.4% of the manual process average counts. To evaluate manual and robotic G-Rex mixing, primary human T cells in two volumes (100 versus 600 mL) and a range of cell concentrations (0.08x10^6^ - 13.7x10^6^ cells/mL) were placed into a G-Rex 100M-CS bioreactor (Figure 4g, h). The cells were allowed to settle for 2 hours and then resuspended manually or robotically with a swirling motion. Cells were then sampled and counted. Averaged manual and robotic cell counts from 3 replicates across four conditions showed comparable results between manual and robotic methods, with less than 2% difference.

### 2.5 Cell Counting

Cell counting and viability assessment is performed at many points of the cell therapy manufacturing process. Cell counting is manually performed by drawing a small volume of liquid (cell suspension) from a bioreactor or cell culture bag. The liquid is loaded into a cassette which is inserted into a cell counting instrument. We mimicked these manual process steps robotically (Figure 5a-c). The cell counting module uses an automated NucleoCounter NC-200 instrument with a custom designed cartridge to house three robotic cassettes (Figure 5d,e). Each robotic cassette has a fluidic chamber where the cell suspension is dispensed and temporarily stored. Then a cell counting cassette containing immobilized staining solution is dipped in the fluidic chamber and cell suspension is drawn in the cassette. To start a cell counting protocol on the NC-200, the robot uses a custom end-effector which allows it to press the "RUN" button on the instrument. Once the cell count is complete, the software acquires the cell count files from the counter using the SMB file sharing communication protocol. These results are then stored and shared with the operators within the digital record as soon as they are retrieved.

**Figure 5:**
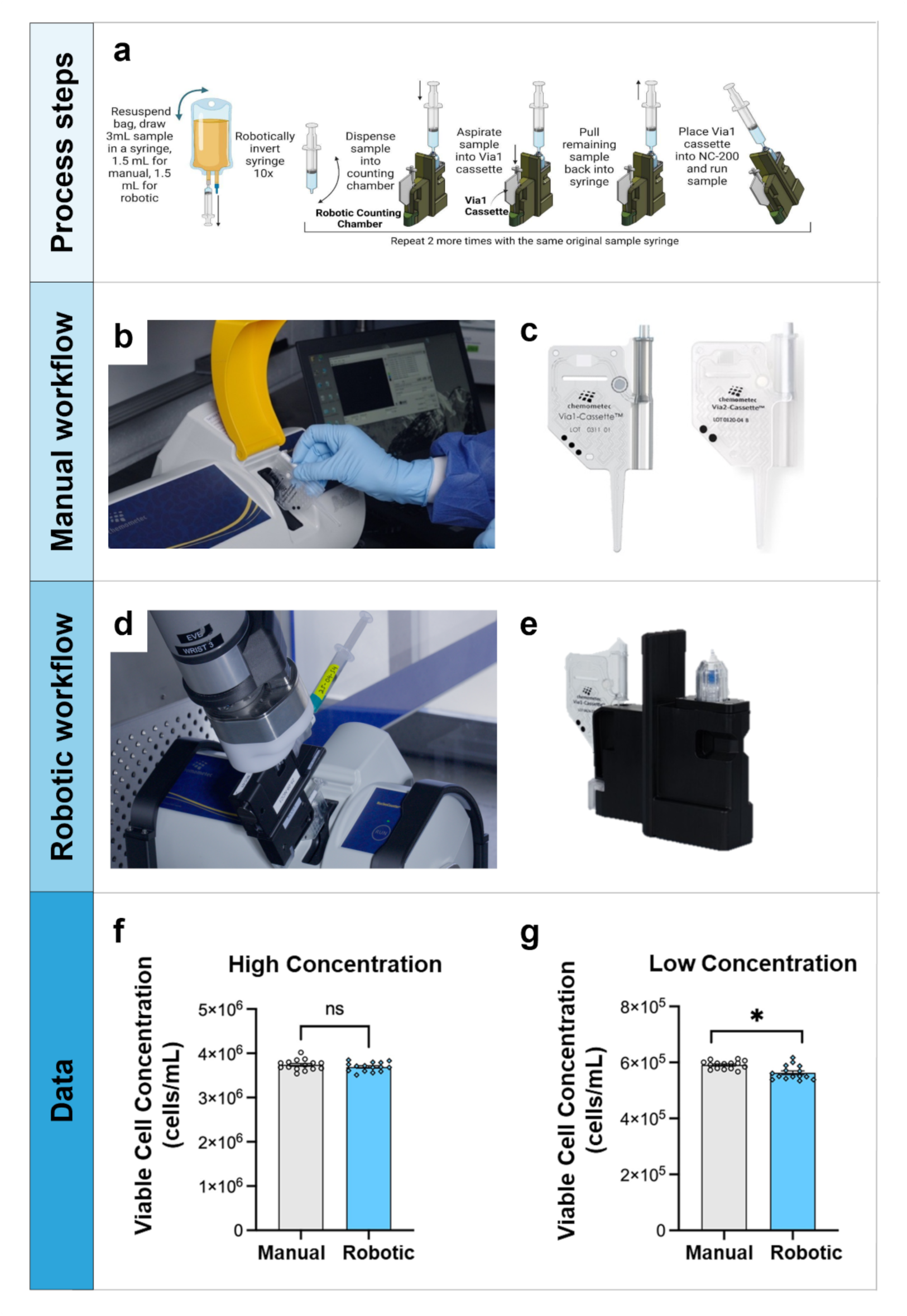
Comparison of manual versus robotic cell counts at high and low concentrations using a NucleoCounter NC-200 cell counter. (a) Schematics showing robotic cell counting workflow. (b-c) Image of operator performing cell count (b) and off-the-shelf Via-1 cassette (c) used in manual workflow. (d-e) Image showing robotic arm inserting the robotic cassette in NucleoCounter NC-200 for cell count. (f-g) Data showing manual and robotic comparison of viable cell concentration at high concentration (left) and low concentration (right) (unpaired t-test, for each concentration, n = 15 cell counts for manual and n = 15 cell counts for robotic). * indicates statistically significant results, p < 0.05.

To compare side-by-side the manual and robotic cell counting, human T cells were placed in bags at two concentrations: 3.5 - 4.0 x10^6^ cells/mL (“high”) and l5.3 - 6.1 x10^5^ cells/mL (“low”). Using 10mL syringes, a small volume of cell suspension (∼3mL) was drawn from the bag and cells were counted on a NucleoCounter NC-200 by either a manual operator or a robotic process. The cell concentrations measured were comparable between manual and robotic approaches. For the high concentration, the average result was 3.74x10^6^cells/mL (manual, n = 15) and 3.69x10^6^cells/mL (robotic, n = 15). The coefficients of variation (CV) were similar: 3.23% (manual) and 2.77% (robotic). Results were also comparable at the low concentration although a statistically significant lower count for the robotic condition was observed: 5.91x10^5^ cells/mL (manual, n = 15) and 5.63 x10^5^ cells/mL (robotic, n = 15. The CV was also comparable: 2.41% (manual) and 4.47% (robotic) (Figure 5f).

### 2.6 Cell Selection

Cell selection or isolation is performed to obtain a purified population of target cells, such as T cells, from the heterogeneous cell population present in blood. In the robotic system, we incorporated a Cytosinct 1000 automated cell selection platform. The instrument is designed to isolate specific cell types from a heterogeneous cell population using an immuno-magnetic cell separation process (Figure 6a). The key components of the instrument include a liquid sensor to monitor the entry of sample into the consumable kit, a magnetic separation unit, a peristaltic pump to control the fluid flow rate, and pinch valve to control flow direction within the kit (Figure 6b). Proper integration and positioning of the components present on the consumable kit onto the instrument are essential for successful cell separation. The cell selection module was automated to meet the following requirements: the instrument uses a standard consumable, the module autonomously executes protocols, and the performance is equivalent to a system run manually.

**Figure 6.**
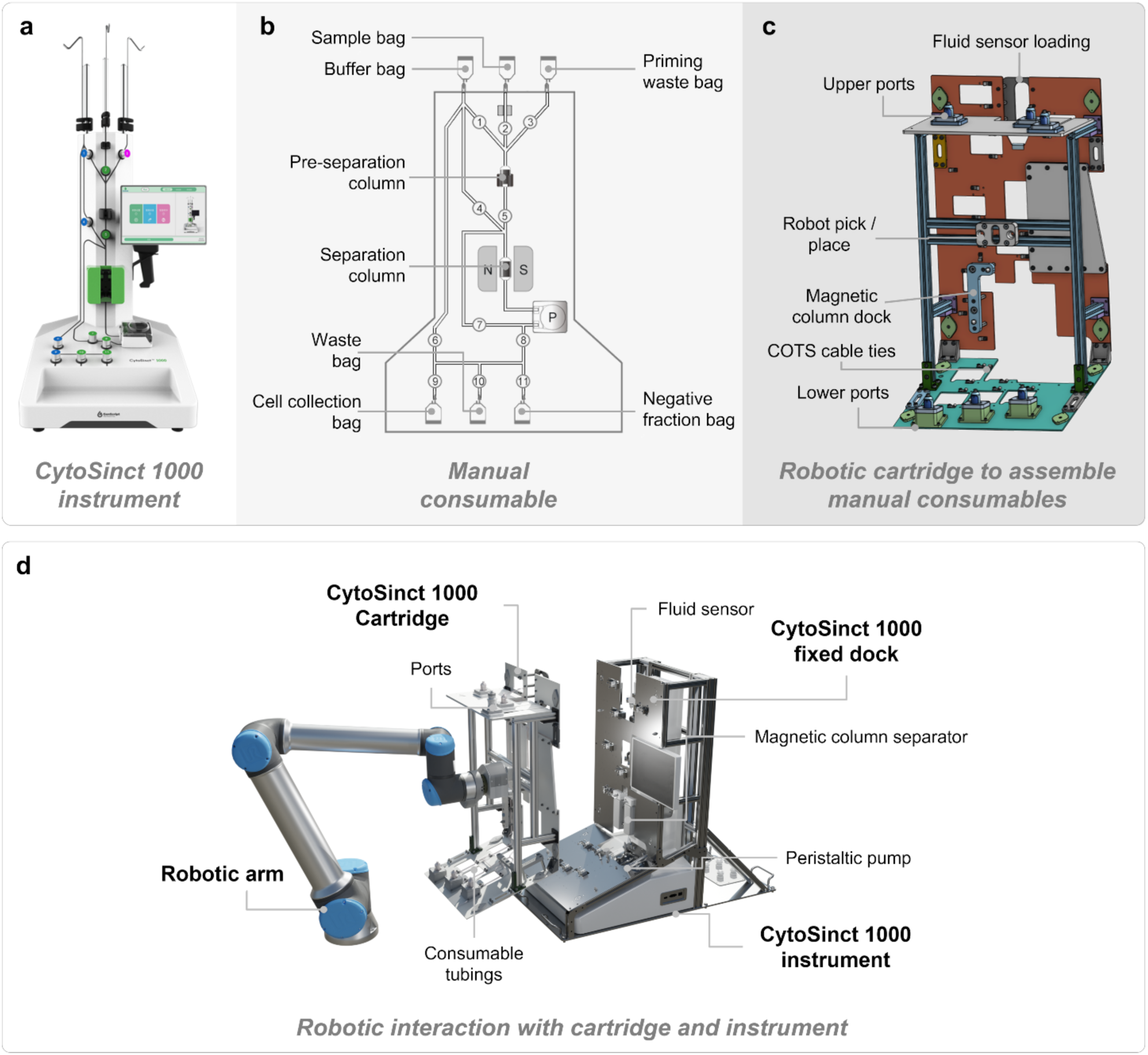
Automation strategy for CytoSinct 1000 magnetic cell separation instrument. a) Image of off-the-shelf instrument used in manual workflow. b) off-the-shelf standard consumable used in manual cell selection workflow. c) Cartridge to assemble standard consumable for automated process d) 3D render showing different components for instrument automation: robotic arm interacting with robotic cartridge and off-the-shelf instrument with dock plate to house robotic cartridge.

The robotic Cytosinct consumable kit was added onto the instrument utilizing the Cytosinct 1000 cartridge (Figure 6c). There were two major challenges associated with the consumable loading task. First, the cartridge has a tall profile, rendering accurate pick/place difficult due to risk of cantilevering and structural deflection. To address this challenge, the robotic pick/place interface is located at the cartridge’s center of mass (Figure 6c). Both planes of the cartridge were roughly aligned with the Cytosinct 1000 fixed dock (Figure 6d) using slots that mate with alignment pins on the dock. After loading, electromagnets are engaged to hold the cartridge in place. The second load challenge was that the Cytosinct 1000 instrument has eleven different pinch valves requiring lateral tube insertion. An augmented valve approach was thus developed to support front-loading of the tubes. A break-beam sensor detected whether the instrument’s valve was closing, and the augmented valve closure followed within milliseconds. To communicate with the instrument without requiring any usage of the touchscreen by the operator, a custom API was developed in collaboration with the manufacturer, GenScript. Specifically, the software can communicate with this API using the TCP/IP communication protocol, allowing the operators to bypass any steps requiring the touchscreen on the instrument and run any custom protocol.

Cell selection on the CytoSinct is performed by labelling a heterogeneous cell suspension with magnetic beads that bind specifically to T cells. The labelled cells are then processed on the CytoSinct 1000 instruments using the TS consumable kit (Figure 7a). The cell selection module (Figure 1b) contains a CytoSinct 1000 magnetic cell separation system and bag stations (Figure 7b). An off-the-shelf CytoSinct 1000 Tubing Set was modified by adding tubing with a SmartSite port to enable closed system connection with the bags and assembled on the robotic cartridge (Figure 7c). The relevant bags were custom designed to be compatible with the robotic loading or unloading onto the enrichment/isolation module bag stations. The robotic cartridge and bags were introduced into the robotic system via the input/output module and the robotic arm shuttled it to the enrichment/isolation module (Figure 7d). Once the robotic cartridge with tubing kit is docked and all the key components are robotically placed on the instrument, the relevant bags are connected to the tubing kit and cell selection protocol is initiated (Figure 7e).

**Figure 7.**
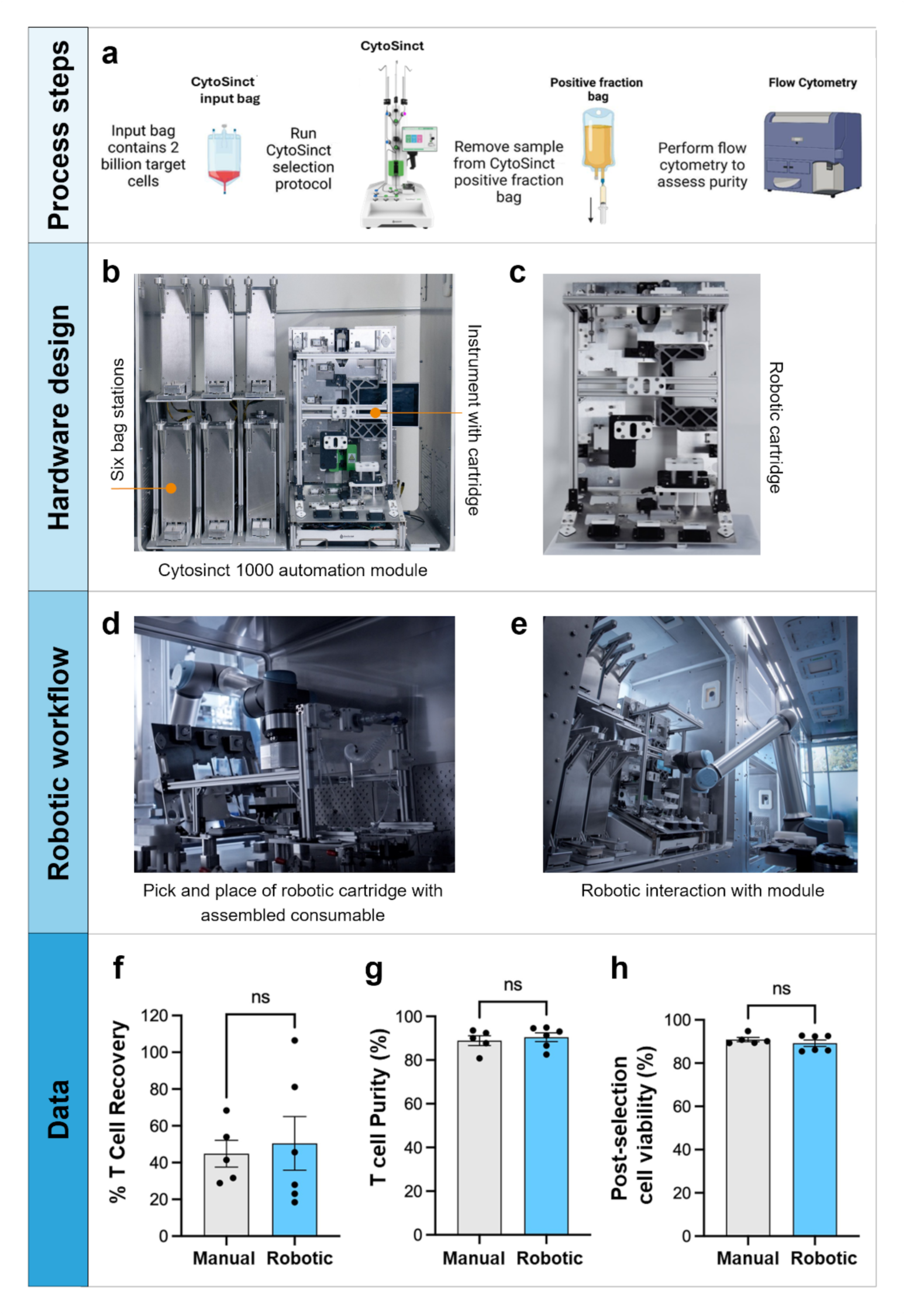
T cell recovery and purity comparison between manual and robotic cell selection using a CytoSinct 1000 magnetic cell separation system. (a) Schematics of the robotic and manual T cell selection workflow. (b) Image showing the CytoSinct module housing the automated instrument with cartridge assembled and six bag stations (c) Image of the robotic cartridge on which off-the-shelf consumable is assembled. (d) Image showing robotic arm picking up robotic cartridge with consumable assembled from input/output module (e) Robotic arm interacting with the CytoSinct module. (f) Percent recovery of T cells in the positive fraction compared to the starting material, g) post-selection T cell purity in the positive fraction, and (h) post-selection cell viability. Statistical comparisons are an unpaired t-test, ns = non-significant. n = 5 for manual condition and n = 6 for robotic condition.

We performed a concurrent manual (n = 5) and robotic (n = 6) positive selection of T cells using a CytoSinct instrument and the CytoSinct 1000 TS tubing kit. Frozen leukapheresis containing heterogeneous cells was thawed, washed, and incubated with CD4 and CD8 magnetic microbeads. The target cell input range was 0.5x10^9^- 2.0x10^9^ cells. Post-selection, the total T cell yield recovery was comparable between manual and robotic conditions, although greater variability was observed in the robotic condition compared to manual (Figure 7f). Post-selection T cell purity was assessed by flow cytometry (Figure 7g). The percentage of CD3+ T cells in the positive fraction was comparable between conditions, and above 80% in all runs. Post-selection cell viability was quantified in the positive fraction for both the manual and robotic processes (Figure 7h). Comparable results for cell viability were observed in both processes. Other cell subsets were also quantified (Supplementary Material, Figure S1).

### 2.7 Buffer Exchange and Cell Harvest

The CTS Rotea is a closed automated counterflow centrifugation system that has several functions, including cell separation based on size and density, volume reduction, cell wash, and buffer/media exchange (Supplementary Material, Figure S2a). To perform these functions, a closed single-use sterile consumable kit (Supplementary Material, Figure S2b) is used. The instrument is composed of several key components: a peristaltic pump for fluid movement, valves to direct fluid flow, a centrifuge cone for density-based separation of fluid contents, and various pressure and bubble sensors that act as step triggers and aid in troubleshooting. Before operating the instrument, various bags are welded to the consumable kit’s tubing. Different protocols are then executed on the instrument, depending on the specific application.

The CTS Rotea consumable kit was mounted onto the instrument using a custom aluminum cartridge, designed to enable robust, repeatable handling by the robotic arm. The cartridge was seated onto the fixed dock on the instrument via locating pins, then secured in place using two cams, one on each side of the instrument (Supplementary Material, Figure S2c,d). Following the large-scale docking of the full consumable cartridge onto the Rotea platform, several smaller, precise loading steps were required to complete setup. The pump load mechanism (Supplementary Material, Figure S2d) uses a specific pump motor to transcribe a circular arc to fully seat the tubing in the peristaltic pump. The centrifuge cone was placed into its chamber using an integrated alignment feature on the CTS Rotea cartridge itself, ensuring precise positioning. For centrifuge cone removal, a photoelectric sensor on the instrument’s centrifuge chamber clip detected the cone’s orientation, and the cone included a dedicated robotic gripper interface for independent extraction based on orientation. To ease insertion of each tube into its respective bubble sensor, a pre-load compression jig was used to stretch the tubing, reducing its diameter for smoother engagement with each bubble sensor’s clamshell-shaped tube housing. Foam inserts were added between the centrifuge cone and the robotic cartridge to eliminate rattling during high-speed centrifugation steps.

To confirm the performance of the automated Rotea, we performed cell washing and buffer exchange both manually and using an automated (robotic) process. Cells were harvested into a 1L bag, and both the input cell bag and reagent bags were loaded into either the manual or robotic Rotea system. Cell recovery and viability were assessed by performing cell counts before and after the Rotea process (Figure 8a). For the robotic process, the counterflow centrifugation module (Figure 1b) consists of bag stations, a robotic cartridge for holding the modified consumable kit and a dock for holding the Rotea instrument (Figure 8b,c). The consumable kit is first loaded into the input/output module, then transferred to the centrifugation module using a robotic cartridge gripper (Figures 8d, e). Once correctly installed, the fluid bags are automatically connected to the consumable kit positioned on the instrument. We compared the cell recovery and manual versus robotic buffer exchange using the Rotea and primary human T cells (ranging from 0.2x10^9^ to 4.0x10^9^) in independent runs. The cell recovery for the manual process ranged from 83% to 95%, while the cell recovery for the robotic process ranged from 81% to 133% (Figure 8f). The cell viability post-harvest was above 93% in the manual condition, whereas in the robotic condition, viability ranged from 83% to 97% (Figure 8g). There was no significant difference in percent cell recovery or post-harvest cell viability between manual and robotic conditions.

**Figure 8:**
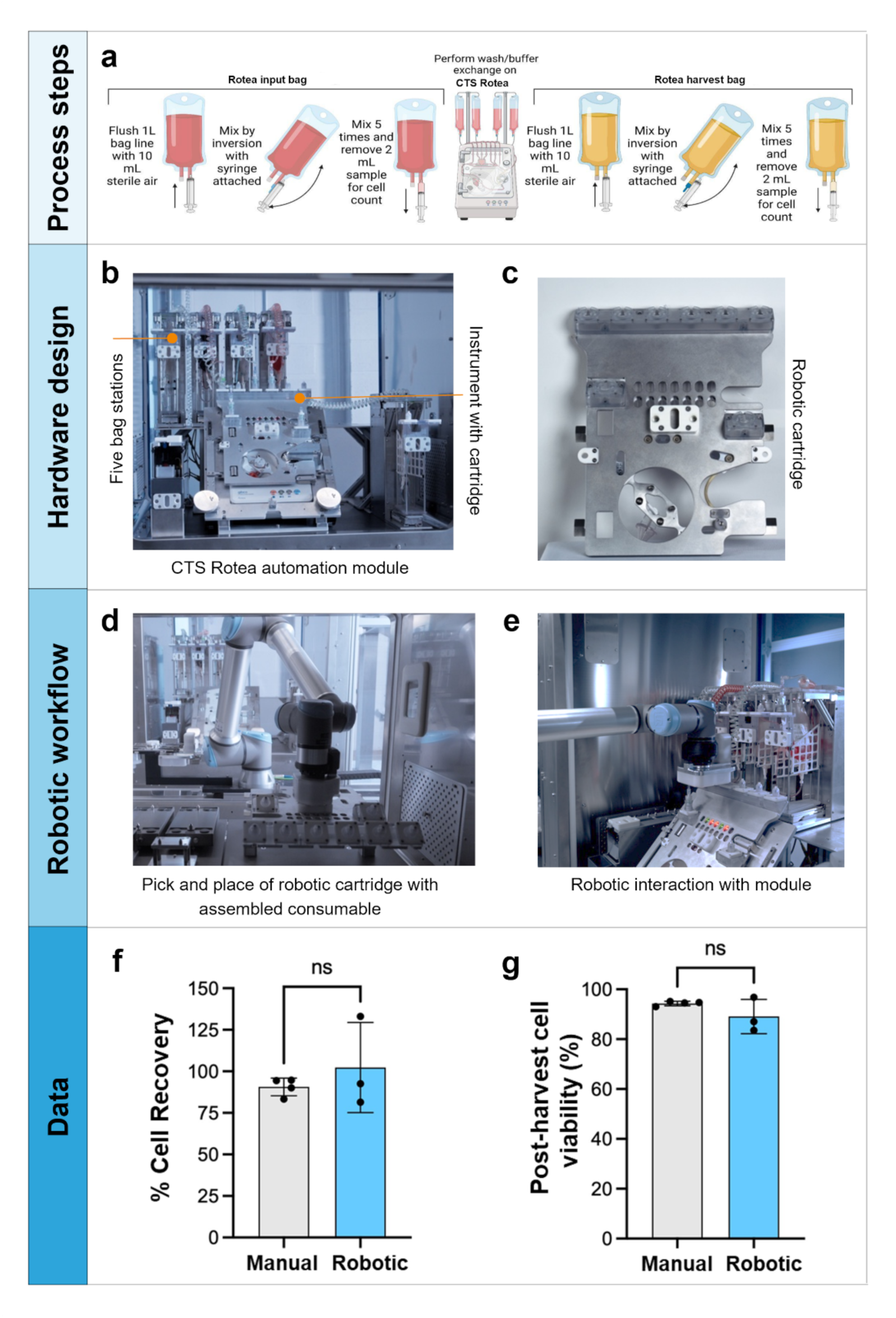
Comparison of Manual and Robotic Harvest Processes Using the CTS Rotea Counterflow Centrifugation System. (a) Schematics of the robotic and manual harvest workflow. (b) Image showing the CTS Rotea module housing the automated instrument with cartridge assembled and five bag stations (c) Image of the robotic cartridge on which off-the-shelf consumable is assembled. (d) Image showing robotic arm picking up robotic cartridge with consumable assembled from input/output module (e) Robotic arm interacting with the CTS Rotea module. (f) Data showing percentage cell recovery, and (g) post-harvest total cell viability between manual and robotic T cell harvest (unpaired student’s t-test, non-significant difference between manual and robotic condition. n = 4 for manual and n = 3 for robotic).

### 2.8 Aseptic Process Simulation (Media Fill)

To confirm the ability of the robotic system to maintain sterility during cell manipulations, aseptic process simulations were carried out for a model CAR-T cell manufacturing process. In the media fill experiment, the unit operations were performed sequentially and included transfer of volume between syringe and G-Rex bioreactor, transfer of volume from bag to G-Rex bioreactor, and resuspension of G-Rex bioreactor, overall representing over 350 connections. No bacterial growth was observed in the media after 14 days of incubation at 37 °C or longer (Supplementary Material, Table S2). Notably, the robotic system was found to maintain sterility of the process when placed in a standard research laboratory space (non-classified environment).

## 3. Discussion

We describe a modular robotic system that performs several common cell therapy manufacturing operations with comparable results to a trained human operator. The robotic system effectively replicated complex manual processes using the same standard equipment and consumables. Importantly, the robot can mimic human motions when required for a process. For example, resuspension of cells in a G-Rex 100M CS bioreactor was based on motion capture from a human operator. Optimizing motions for each specific process is a key area of potential improvement. In some cases, software changes to instruments were necessary to allow for automation. For example, to improve cell recovery in the robotic runs, we made changes to the Enrichment 1.1 protocol of the CytoSinct such as increasing the washing/elution time. Similar optimizations are likely to be required for instruments with large, dedicated tubing sets.

There are still several important processes that were not automated. For example, we did not demonstrate the precision and accuracy of partial large volume transfer (>100 mL) and cell transfer efficiency from one vessel to another. These improvements are planned for further development. In addition, small volumes (<1 mL) are also challenging for the robot. For small volumes, careful calibration must take into account viscosity, surface tension, dead volumes, and consumable size.

Importantly, all operations need to be done under conditions that maintain sterility of the product. The robotic system uses multiple-use needle free connectors, which were sanitized with alcohol-containing disinfecting caps before each use. Since all activities occur inside of the robotic cluster without human intervention, the risk of culture contamination is likely to be reduced compared to manual processes because humans are the primary source of microbial contamination in a cleanroom. The absence of human operators may enable a lower cleanroom classification (Grade C instead of Grade B) for the interior of the robotic cluster. For use in GMP manufacturing processes, the interior of the cluster will require regular cleaning and environmental monitoring. However, empirical data is required to determine the scope of such cleaning and monitoring.

Advanced architectures incorporating multi-arm parallelism and concurrent processing of multiple patient batches will significantly increase throughput and operational scalability. As the robotic actions become more reliable, their processing rate will be increased, thus decreasing the overall time of operation. Future iterations will allow users to customize process parameters, modify workflows mid-run, and integrate diverse vessel types and consumables, enabling adaptation across a wide range of cell therapy modalities. In parallel, standardization, optimization, and cost-efficient mass production of the robotic adapters for each consumable and piece of equipment is required to be able to scale out the system to commercially relevant numbers. Modules enabling automated handling of a variety of consumables are under development.

Furthermore, to address full end-to-end automation, new instruments will be automated to perform immunolabelling, small volume harvest, product formulation, and fill-finish operations. Future systems will integrate robust fault-tolerant control algorithms, automated error detection and recovery mechanism, real-time barcode-based traceability for consumables, reagents and samples. The incorporation of imitation learning and reinforcement learning frameworks will facilitate anthropomorphic motion planning, improving task adaptability, and reducing training overhead. Furthermore, digital twin modeling and simulation will enable predictive validation of robotic workflows, optimizing feasibility assessments prior to live execution.

## 4. Conclusions

Robotic automation has the potential to perform many manual procedures in cell therapy manufacturing. End-to-end automation of a modular manufacturing process may increase the supply of cell therapy products to meet global demand, improve efficiency, accelerate delivery timelines, and ensure high quality standards are consistently met.

## 5. Materials and Methods

The robotic workflow for each unit operation is shown in the Supplementary material, Movie S1.

### 5.1 Smart-site and Texium connectors

The robotic connectors are based on commercially available Luer connectors, with the addition of external components that increase the robustness of the automated connection/disconnection task (Figure 2). The standard connectors used during the work are the Beckton-Dickson SmartSite (closed female Luer) and the Beckton-Dickson Texium (closed male Luer). The SmartSite and Texium are closed, needle-free connectors with Luer threads. Both connectors are equipped with a membrane that keeps their default configuration “closed”, preventing any flow through the connector and sealing the internal environment from the outside. The membrane is pushed away into the “open” configuration when a matching Luer lock component is connected, and returns to the original “closed” state when the Luer lock component is disconnected. The SmartSite and Texium connectors are classified as Class II, Sterile, Closed System Transfer Devices (CSTD) and approved by the FDA for use in clinical/hospital settings (re: K223088 for the SmartSite, and K223076 for the Texium). Needle-free connectors similar to the SmartSite are currently used on several GMP-proven instruments that are employed in the commercial manufacturing of FDA-approved cell therapies, including the G-Rex bioreactor (by Wilson Wolf). For disinfecting SmartSite and Texium, we utilized the Merit Medical DualCap Disinfection & Protection System.The DualCap Disinfection & Protection System is FDA approved (re: K142806).

### 5.2 Vial-to-Syringe Volume Draw

Each vial for robotic manipulation was attached to a 20 mm vented Vialok (Yukon) and SmartSite (Becton Dickinson) and then assembled to a proprietary vial adapter. Each vial was filled with 5 mL dyed water. For robotic conditions, the two filled vials were placed in the in-house designed vial cartridge. An empty 10 mL syringe with Texium connectors was attached to the coupler and placed in the designed syringe cartridge (Figure 3a). 4 mL was drawn from vial 1 and 1.8 mL was drawn from vial 2 using the same syringe to reach a target volume of 5.8 mL. Similarly, in the manual condition, 5.8 mL of fluid from two vials was pulled using the 10 mL syringe attached to the connector. The weight of the syringe for both conditions were measured manually and collected pre-draw (tare weight of the empty syringe), post-draw from vial 1, and again post-draw from vial 2 (Analytical Balance, Ohaus).

### 5.3 Bag-to-Bag Cell Transfer

Human T cells (StemCell Technologies & AllCells) were seeded into an automation compatible permalife bag (OriGen, PL120-2G) in 100 mL TexMACS media. 24 hours post-seeding, cells were activated using TransAct (Miltenyi, 130-111-160). On day 3, cells were transferred from the permalife bag to the transfer pack (Fenwal Inc, 4R2001) robotically and the percentage of cells in each bag was assessed (Figure 3b).

### 5.4 Bag/Bioreactor Resuspension and Sampling

Robotically enriched and expanded human T cells were harvested on day 8 post-activation from G-Rex 100M-CS (Wilson Wolf Manufacturing Corporation, J81100-CS) into a 1L transfer bag (Terumo, BB*T100BB71), resuspended, and sampled for cell counting (Figure 4a). Post-harvest, the cells in the bag were subjected to buffer exchange using the robotic CTS Rotea counterflow centrifugation system. The washed cells were then collected in another 1L transfer bag, resuspended, and sampled for cell counting. To accomplish robotic bag resuspension and sampling, the robot attached a syringe filled with 10 mL of sterile air to the 1L transfer bag to flush the tube line. Next, the bag was rocked for 2 minutes to mix the cells by tilting it up and down at a 30 degree angle from the horizontal position, with the syringe still attached. Following resuspension, the robot mixed the bag line with the syringe five times. A 2 mL sample was then removed from the bag using the same syringe. The sample and bag were then output from the robotic system and cell count was performed. The same 1L bag was then manually rocked for 2 mins by tilting it up and down at a 30 degree angle from the horizontal position, and a 2 mL sample was manually removed via syringe. All cell counts were performed manually using the NucleoCounter NC-202. This process was repeated on six different 1L transfer bags, with resuspension and sampling after G-Rex 100M-CS transfer (n = 4) or post-Rotea (n = 2).

For the bioreactor resuspension operation, expanded primary human T cells were divided into four experimental conditions in different G-Rex 100M-CS bioreactors, which included variations in cell number and volume: low cell number, low volume (5.3x10^7^ cells in 100 mL medium), high cell number, high volume (1.9x10^9^ cells in 600 mL medium), low cell number, high volume (4.9x10^7^ cells in 600 mL medium), and high cell number, low volume (1.3x10^9^ cells in 100 mL medium) (Figure 4b). The robotic cell resuspension protocol was designed to replicate the manual resuspension motion: a scientist from Stanford’s Laboratory for Cell and Gene Medicine performed multiple resuspensions with motions recorded and translated to the robotic system using a motion capture system (Opitrak). After each resuspension, all conditions were allowed to rest for 2 hours to allow the cells to settle before the next resuspension.

### 5.5 Cell Counting

T cells were expanded and transferred into a bag at different concentrations. A 3.5 mL sample was drawn from the bag using a 10 mL syringe (Becton Dickson) by either the robotic arm or a human operator (Figure 5a). The cells were loaded into a Via1-Cassette (Chemometec, 941-0011). The robotic process used a specialized reservoir to perform the cell transfer from syringe to the counting cassette. Cell counts were performed in triplicate either manually or robotically using NucleoCounter NC-200.

### 5.6 Cell Selection

Frozen or fresh apheresis was washed with phosphate buffered saline/EDTA (Miltenyi) containing 0.5% human serum albumin and incubated with CD4 and CD8 microbeads (Miltenyi) for 30 minutes using LOVO Cell Processing System (Fresenius Kabi). The labelled cells were equally divided into manual and robotic conditions for cell enrichment and isolation using the CytoSinct 1000 magnetic cell separation system. TexMACS media supplemented with 3% serum, IL-7 and IL-15 was used as the buffer for enrichment. For manual conditions, the CytoSinct 1000 TS kit (GenScript Biotech Corporation, D00029) was set up per manufacturer’s instructions. For robotic conditions, the kit and bags were assembled onto proprietary cartridges and loaded to the robotic cluster. For both manual and robotic conditions, 2x10^9^ target cells were loaded and Enrichment 1.1 protocol was performed. The cell count and viability pre-enrichment were provided by Stanford (Cellometer Auto 2000, Nexcelom Bioscience). Post-enrichment counts were assessed using NucleoCounter NC-202. To quantify T cell purity in the positive fraction, CD3+ T cells were measured by flow cytometry (BD LSR Fortessa, BD Biosciences or CytoFlexLX, Beckman Coulter) performed either at University of California San Francisco (UCSF) flow core or at Stanford University flow core. For each run, separate healthy donor apheresis was used (Figure 7a).

### 5.7 Buffer Exchange and Cell Harvest

Enriched T cells were expanded and subjected to wash/buffer exchange using the CTS Rotea Counterflow centrifugation system and CTS Rotea Single-Use Kit (Thermo Fisher Scientific, A49585) (Figure 8a). The published buffer exchange protocol from Thermo Fisher Scientific was performed in both the manual and robotic conditions. However, prime volumes were increased in the robotic condition to account for the additional tubing length. Plasmalyte A containing 4% HSA was used as the buffer for both conditions. Cells were harvested in 40 mL of buffer.

### 5.8 Aseptic Process Simulation (Media Fill)

Sterile Tryptic Soy Broth media (Corning, MT46060CM) was transferred into pre-sterilized 1L transfer bags and containers needed for the process simulation in an ISO 5 biosafety cabinet (BSC). The prepared materials were then transferred into the robotic cluster via the input/output module. Here the media was robotically transferred between syringe and G-Rex 100M-CS bioreactor, from bag to G-Rex bioreactor, and resuspended in the latter. At the end, containers (n = 4) were filled with the processed media and checked for sterility in accordance with USP. Post 14-day incubation media growth promotion testing was also performed. To challenge the closed connection capability of the robotic clusters, the above aseptic process simulation was carried out under the worst possible conditions, *i.e.* having the unit self-contained HEPA/HVAC system purposefully non-operational.

### 5.9. Statistical Analysis

Graphpad Prism 10.4.0. were used to perform statistical analysis. Results are shown as mean ± Standard Deviation (SD) or mean ± Standard Error of Mean (SEM) and statistical differences between manual and robotic conditions were assessed by paired or unpaired Student’s t-test. A two-tailed p value < 0.05 indicates a significant difference.

## Acknowledgement

We would like to express our gratitude to ChemoMetec, Akron Bio and Cytiva, for technical advice and feedback during the work.

## Funding

This work was supported in part through a collaboration between Stanford Center for Cancer Cell Therapy and Multiply Labs, Inc.

**Movie S1:** <u>Unit Operations Paper Videos</u>

---

**Figure S1:**
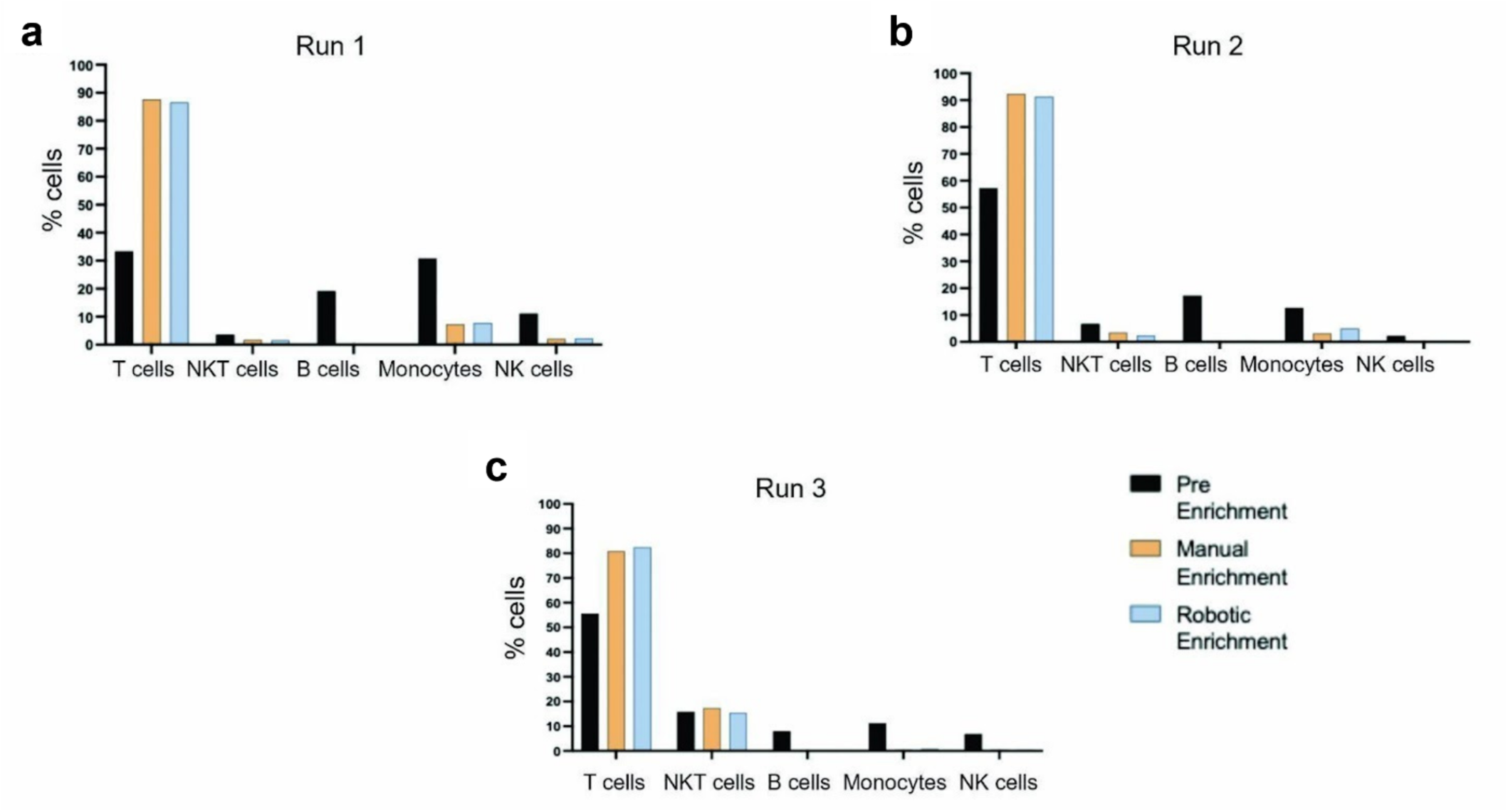
Percentage of different cell populations post-selection were assessed in the positive fraction after CD4 and CD8 cell selection on CytoSinct 1000 using flow cytometry (n = 3).

**Figure S2.**
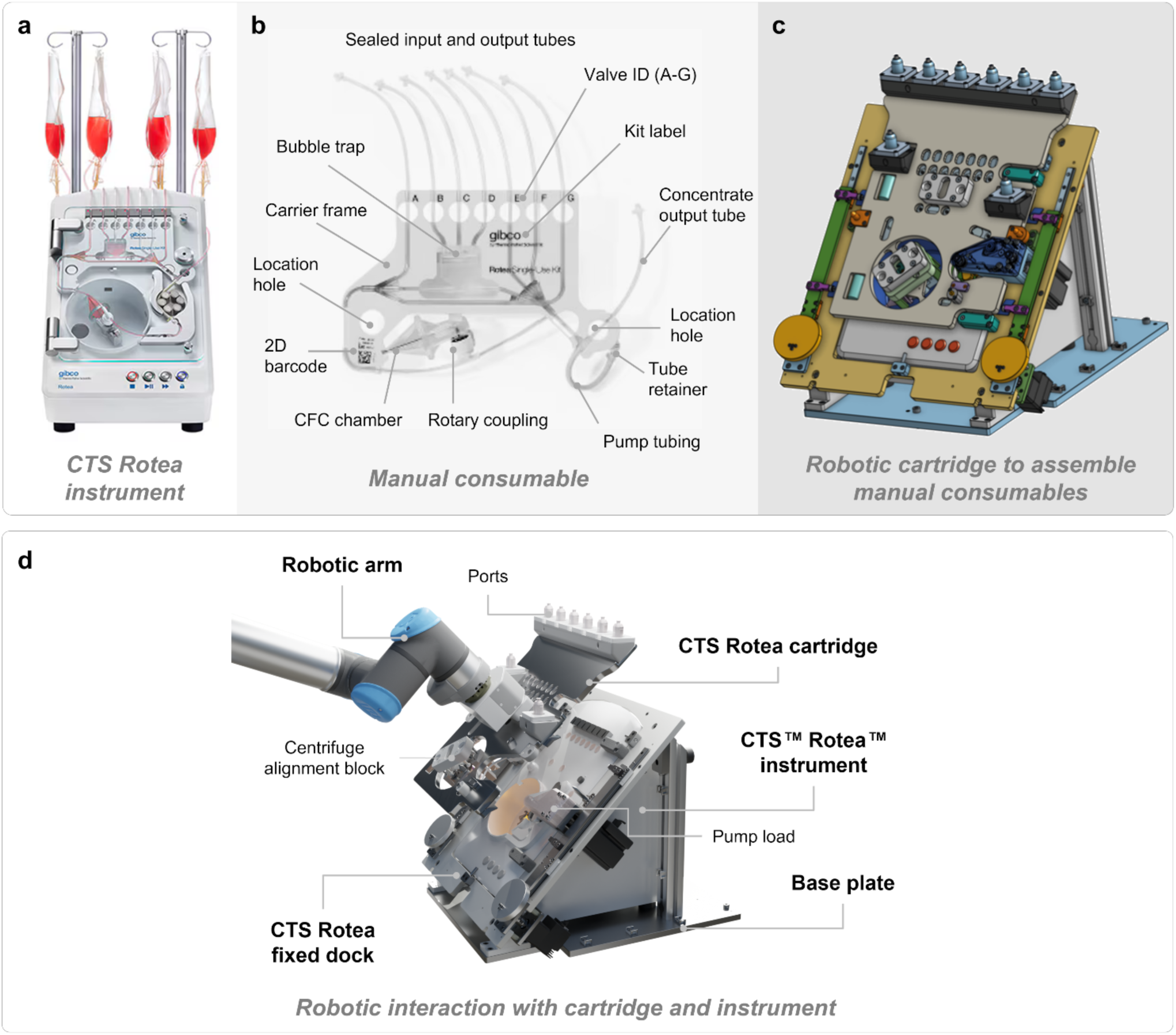
Automation strategy for CTS Rotea counterflow centrifugation instrument. a) Image of off-the-shelf instrument used in manual workflow. b) off-the-shelf standard consumable used in manual cell selection workflow. c) Cartridge to assemble standard consumable for automated process D) 3D render showing different components for instrument automation: robotic arm interacting with robotic cartridge and off-the-shelf instrument with dock plate to house robotic cartridge.

**Supplementary Table 1:**
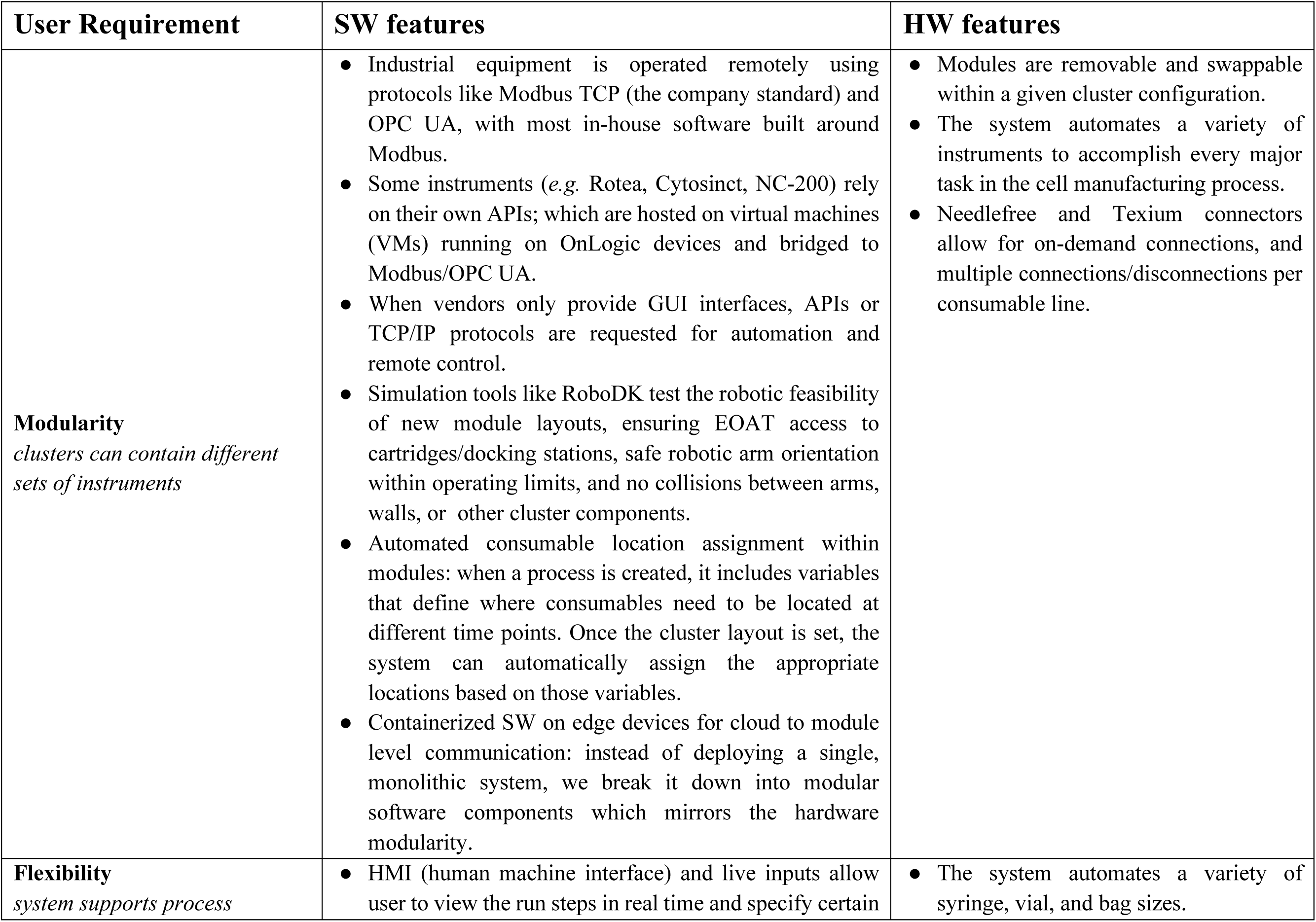
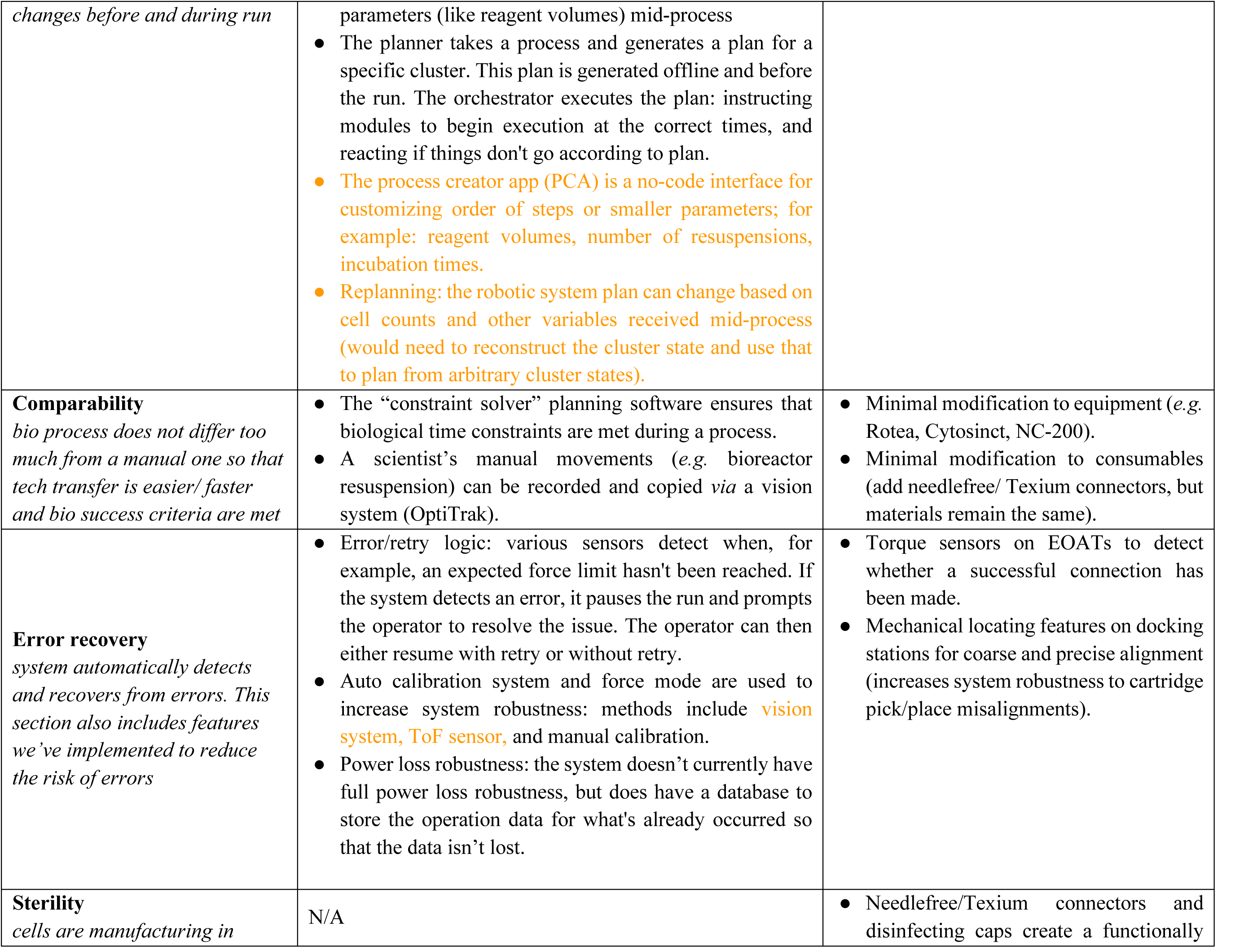
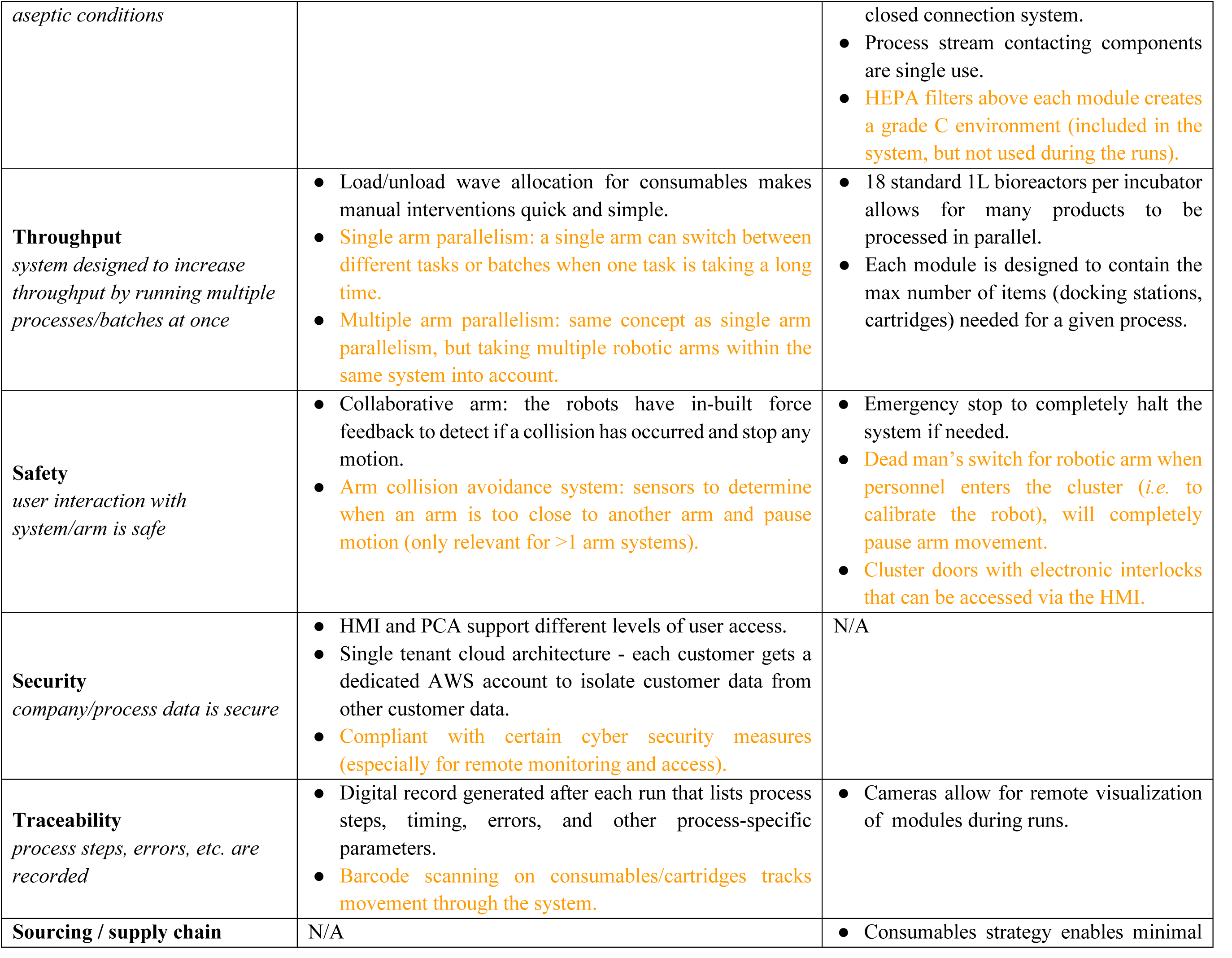
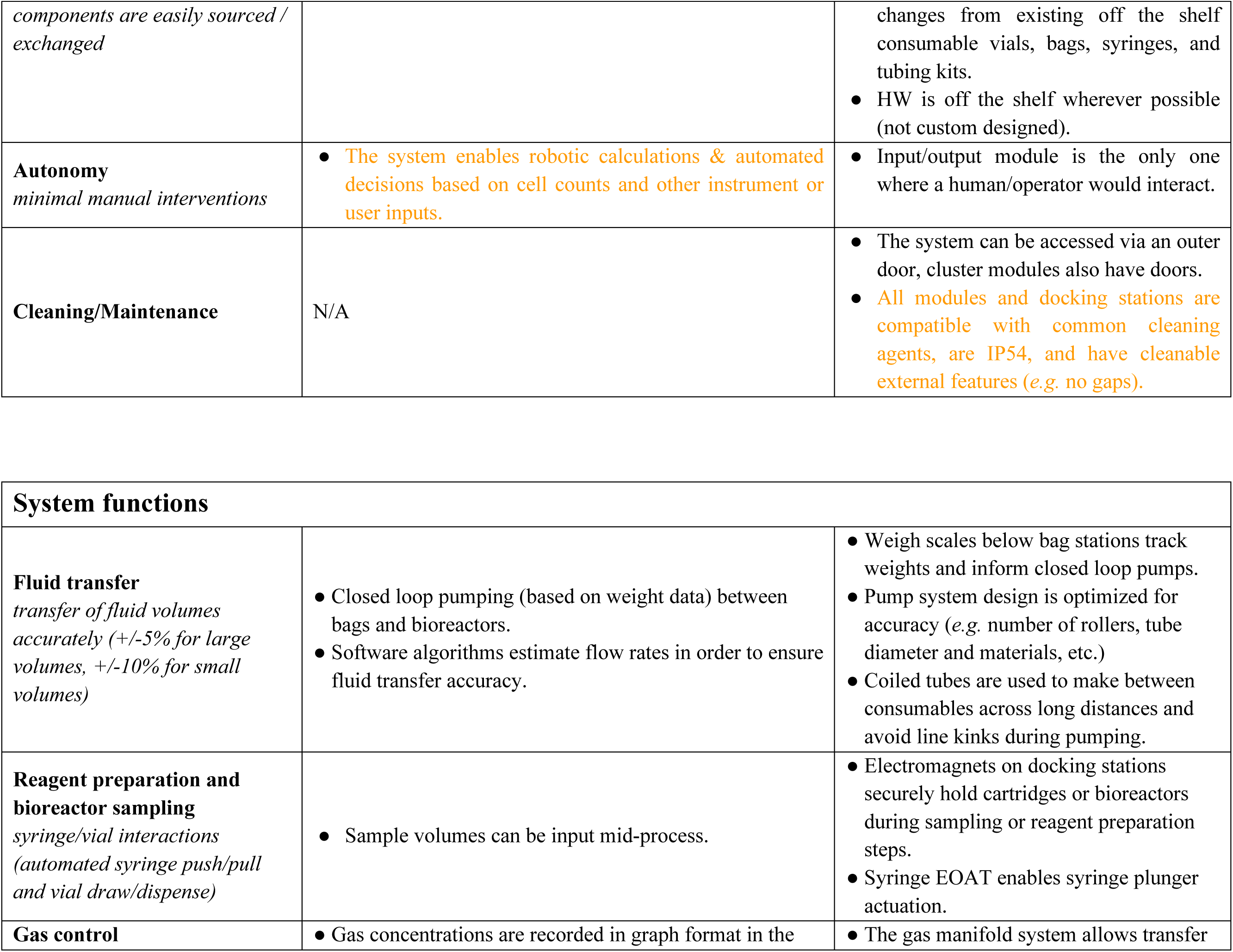
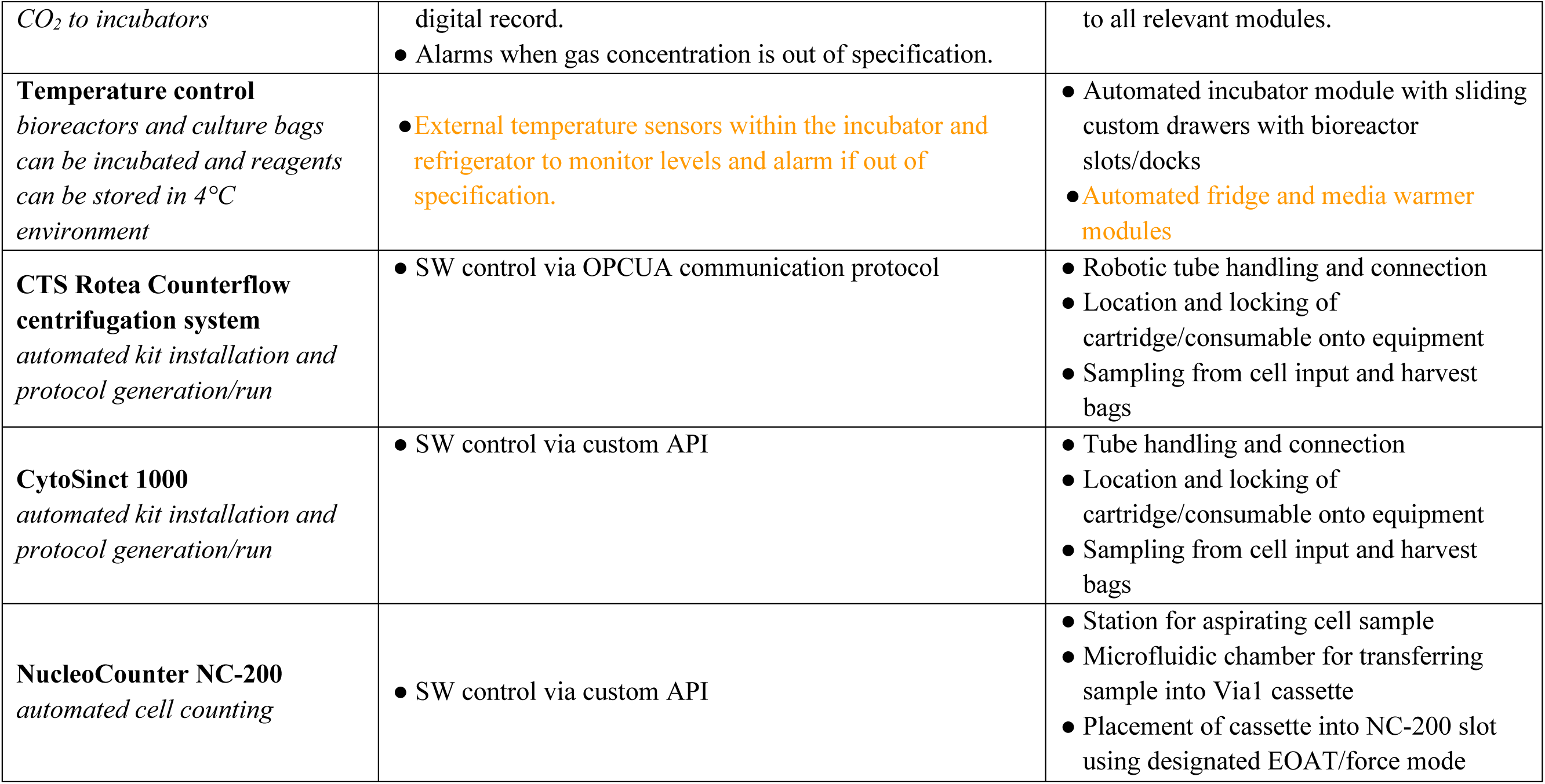
Hardware and software design specifications for the robotic cluster Orange = future/planned improvements (for future iteration of robotic systems)

**Table S2:** Aseptic process simulation results using robotic unit operations for a generic effector T cell manufacturing process.

| Experiment types | Runs | 14-day sterility test<br>(Growth / No Growth) | Pass / Fail |
| --- | --- | --- | --- |
| Media fill | 1 | No growth | Pass |
|  | 2 | No growth | Pass |
|  | 3 | No growth | Pass |
|  | 4 | No growth | Pass |

## Notes

### Competing Interest Statement

-Sudeshna Sadhu, Brigitte Schmittlein, Angela Lares, Christopher Cheng, Xiaojie Chen, Varun
Bhatia, Winston Zha, Sebastien Wha and Sunaina Nayak wish to disclose that they are current
employees of Multiply Labs, Inc. or were employed with the company at the time of this study's
execution. They hold equity in the company;
● Alice Melocchi and Federico Parietti wish to disclose that they are co-founders of Multiply Labs,
Inc. and hold the position of Chief Scientific Officer and Chief Executive Officer, respectively;
● Yifan Yu, Jiawei Wang, Wei Zhang wish to disclose that they are employed by GenScript Biotech
Corporation, which is a company active in the cell therapy manufacturing space;
● Prajakta P. Bhanap and Yongchang Ji wish to disclose that they are employed by Thermo Fisher
Scientific, which is a company active in the cell therapy manufacturing space;
● Andrew Scheffler wishes to disclose that he is employed by ScaleReady, which is a company
active in the cell therapy manufacturing space;
● John Wilson and Dan Welch wish to disclose that they are employed by Wilson Wolf
Manufacturing corporation, which is a company active in the cell therapy manufacturing space;
● Nikolaos Gkitsas-Long, Aidan Jeffrey Retherford, Ramya Tunuguntla and Steven A. Feldman
wish to disclose that they work in the Laboratory for Cell and Gene Therapy Manufacturing
(LCGM) at Stanford University. The LCGM entered a collaboration agreement with Multiply
Labs, Inc. for studying the performance of cell therapy manufacturing robots built by the
company, by comparing them with manual processes;
● Jonathan H. Esensten wishes to disclose he is a paid advisor for Multiply Labs, Inc. He serves
on its scientific advisory board and holds equity in the company. He receives sponsored
research funding from Lonza, Inc. for the development of cellular therapy manufacturing
devices.

### Summary of Updates

The revised manuscript emphasizes the engineering challenges and key design principles underlying various robotic components used to automate standard instruments and consumables in cell therapy manufacturing.

## References

1. De Marco R.C., Monzo H.J., Ojala P.M., 2023, CAR T cell therapy: a versatile living drug, Int. J. Mol. Sci., 24: 6300. 10.3390/ijms24076300

2. Fugger L., Torp Jensen L., Rossjohn J., 2020, Challenges, progress, and prospects of developing therapies to treat autoimmune diseases, Cell, 181: 63–80. 10.1016/j.cell.2020.03.007.

3. Greco R., Alexander T., Del Papa N., Müller F., Saccardi R., Sanchez-Guijo F., Schett G., Sharrack B., Snowden J.A., Tarte K., Onida F., Sánchez-Ortega I., Burman J., Castilla Llorente C., Cervera R., Ciceri F., Doria A., Henes J., Lindsay J., Mackensen A., Muraro P.A., Ricart E., Rovira M., Zuckerman T., Yakoub-Agha I., Farge D., 2024, Innovative cellular therapies for autoimmune diseases: expert-based position statement and clinical practice recommendations from the EBMT practice harmonization and guidelines committee, eClinicalMedicine, 69: 102476. 10.1016/j.eclinm.2024.102476

4. Gustafson M.P., Ligon J.A., Bersenev A., McCann C.D., Shah N.N., Hanley P.J., 2023, Emerging frontiers in immuno- and gene therapy for cancer, Cytotherapy, 25: 20–32. 10.1016/j.jcyt.2022.10.002

5. Park Y., Kwok S.K., 2022, Recent advances in cell therapeutics for systemic autoimmune diseases, Immune Netw., 22: e10. 10.4110/in.2022.22.e10

6. Wang L., Liu G., Zheng L., Long H., Liu Y., 2023, A new era of gene and cell therapy for cancer: a narrative review, Ann Transl Med., 11: 321. 10.21037/atm-22-3882

7. Arabi F., Mansouri V., Ahmadbeigi N., 2022, Gene therapy clinical trials, where do we go? An overview, Biomedicine & Pharmacotherapy, 153: 113324 10.1016/j.biopha.2022.113324.

8. Ginn S.L., Mandwie M., Alexander I.E., Edelstein M., Abedi M.R., 2024, Gene therapy clinical trials worldwide to 2023- An update, J. Gene Med., 26: e3721. 10.1002/jgm.3721

9. https://www.iqvia.com/insights/the-iqvia-institute/reports-and-publications/reports/strengthening-pathways-for-cell-and-gene-therapies, last access on January 22, 2025

10. Ayala Ceja M., Khericha M., Harris C. M., Puig-Saus C., Chen Y. Y., 2024, CAR-T cell manufacturing: Major process parameters and next-generation strategies, J. Exp. Med., 221: e20230903. 10.1084/jem.20230903

11. Heathman T.R., Nienow A.W., McCall M.J., Coopman K., Kara B., Hewitt C.J., 2015, The translation of cell-based therapies: clinical landscape and manufacturing challenges, Regen. Med., 10: 49–64. 10.2217/rme.14.73

12. Iancu E.M., Kandalaft L.E, 2020, Challenges and advantages of cell therapy manufacturing under Good Manufacturing Practices within the hospital setting, Cur. Opin. Biotechnol., 65: 233–241. 10.1016/j.copbio.2020.05.005

13. Campbell A., Brieva T., Raviv L., Rowley J., Niss K., Brandwein H., Oh S., Karnieli O., 2015, Concise review: Process development considerations for cell therapy. Stem Cells Transl. Med., 4: 1155–1163. 10.5966/sctm.2014-0294

14. Lee N.K., Chang J.W., 2024, Manufacturing cell and gene therapies: challenges in clinical translation. Ann. Lab. Med., 44: 314–323. 10.3343/alm.2023.0382

15. Brindley D. A., Wall, I. B., Bure K. E., 2013, Automation of cell therapy biomanufacturing minimizing regulatory risks and maximizing return on investments, BioProcess Int., 11: S42–S49.

16. Doulgkeroglou M. N., Di Nubila A., Niessing B., König N., Schmitt R. H., Damen J., Szilvassy, S. J., Chang W., Csontos L., Louis S., Kugelmeier P., Ronfard V., Bayon Y., Zeugolis D. I., 2020, Automation, monitoring, and standardization of cell product manufacturing, Front. Bioeng. Biotechnol., 8: 811. 10.3389/fbioe.2020.00811

17. Smith D., Heathman T. R. J., Klarer A., LeBlon C., Tada Y., Hampson B., 2019, Towards automated manufacturing for cell therapies, Curr. Hematol. Malig. Rep., 14: 278–285. 10.1007/s11899-019-00522-y

18. Melocchi A., Schmittlein B., Sadhu S., Nayak S., Lares A., Uboldi M., Zema L., di Robilant B., Feldman S.A., Esensten J.H., 2025, Automated manufacturing of cell therapies, J. Control. Release, 381:113561. 10.1016/j.jconrel.2025.02.057

19. Moutsatsou P., Ochs J., Schmitt R. H., Hewitt C. J., Hanga M. P., 2019, Automation in cell and gene therapy manufacturing: from past to future. Biotechnol. Lett., 41: 1245–1253. 10.1007/s10529-019-02732-z

20. Franscini N., Wuertz K., Patocchi-Tenzer I., Durner R., Boos N., Graf-Hausner Ursula U., 2011, Development of a novel automated cell isolation, expansion, and characterization platform, J. Lab. Autom., 16: 204–213. 10.1016/j.jala.2011.01.002

21. Kami D., Watakabe K., Yamazaki-Inoue M., Minami K., Kitani T., Itakura Y., Toyoda M., Sakurai T., Umezawa A., Gojo S., 2013, Large-scale cell production of stem cells for clinical application using the automated cell processing machine. BMC Biotechnol., 13: 102. 10.1186/1472-6750-13-102

22. Kikuchi T., Kino-oka M., Wada M., Kobayashi T., Kato M., Takeda S., Kubo H., Ogawa T., Sunayama H., Tanimoto K., Mizutani M., Shimizu T., Okano T., 2018,. A novel, flexible and automated manufacturing facility for cell-based health care products: Tissue Factory. Regenerative Therapy, 9, 89–99. 10.1016/j.reth.2018.08.004

23. Mantripragada V.P., Luangphakdy V., Hittle B., Powell K., Muschler G.F., 2020, Automated in-process characterization and selection of cell-clones for quality and efficient cell manufacturing, Cytotechnol., 72:615–627. 10.1007/s10616-020-00403-w

24. Murphy M., Barry F., Leschke C., Vaughan B., Gentili C., O’Dea J., Ogourtsov V., Rafiq Q. A., Ochs J., Kulik M., Koenig N., 2017, The autostem platform for closed manufacture of bone marrow-derived mesenchymal stromal cells using a closed, scalable and automated robotic system, Cytotherapy, 19:, s122. 10.1016/j.jcyt.2017.02.199

25. Rafiq Q.A., Twomey K., Kulik M., Leschke C., O’Dea J., Callens S., Gentili C., Barry F. P., Murphy M., 2016, Developing an automated robotic factory for novel stem cell therapy production, Regen. Med., 11: 351–354. 10.2217/rme-2016-0040

26. Thomas R.J., Chandra A., Liu Y., Hourd P.C., Conway P.P., Williams D.J., 2007, Manufacture of a human mesenchymal stem cell population using an automated cell culture platform, Cytotechnol., 55: 31–39. 10.1007/s10616-007-9091-2

27. Fadeyev F.A., Sedneva-Lugovets D.V., Madyarova O.V., 2022, Scale-out cultivation of human dermal fibroblasts using robotic cell culture system: comparison of manual and automated processing, Cell. Ther. Transpl., 11: 63–71, 1 10.18620/ctt-1866-8836-2022-11-2-63-71

28. Herbst L., Groten F., Murphy M., Shaw G., Nießing B., Schmitt R.H., 2023, Automated production at scale of induced pluripotent stem cell-derived mesenchymal stromal cells, chondrocytes and extracellular vehicles: towards real-time release, Processes, 11: 2938, 10.3390/pr11102938

29. Melocchi A., Schmittlein B., Jones A. L., Ainane Y., Rizvi A., Chan D., Dickey E., Pool K., Harsono K., Szymkiewicz D., Scarfogliero U., Bhatia V., Sivanantham A., Kreciglowa N., Hunter A., Gomez M., Tanner A., Uboldi M., Batish A., Balcerek J., Kutova-Stoilova M., Paruthiyil S., Acevedo L. A., Stadnitskiy R.,·Carmichael S., Aulbach H., Hewitt M.,·De Mollerat Du Jeu X., di Robilant B., Parietti F., Esensten J.H, 2024, Development of a robotic cluster for automated and scalable cell therapy manufacturing, Cytotherapy, 26: 1095–1104. 10.1016/j.jcyt.2024.03.010

